# PAC – A novel translational concordance framework identifies preclinical seizure models with highest predictive validity for clinical focal onset seizures

**DOI:** 10.1101/2025.04.04.647239

**Authors:** Lyndsey L. Anderson, Kristopher M. Kahlig, Melissa L. Barker-Haliski, Brian Hannigan, Hamish Toop, Lillian G. Matthews, Jacqueline French, H Steve White, Marcio Souza, Steven Petrou

**Affiliations:** Praxis Precision Medicines, Boston, MA, USA; Department of Pharmaceutics, School of Pharmacy, University of Washington, Seattle, WA, USA; Department of Neurology and Comprehensive Epilepsy Center, NYU Grossman School of Medicine, New York, NY, USA; Center for Epilepsy Drug Discovery, Department of Pharmacy, School of Pharmacy, University of Washington, Seattle, WA, USA

**Keywords:** Focal seizures, Epilepsy, Predictive Validity, Preclinical Models, Translational Concordance

## Abstract

**Objective:** Central to the development of novel antiseizure medications (ASMs) is testing of anticonvulsant activity in preclinical models. While various well-established models exist, their predictive validity across the spectrum of clinical epilepsies has been less clear. We sought to establish the translational concordance of commonly used preclinical models to define models with the highest predictive clinical validity for focal onset seizures (FOS).

**Methods:** The Praxis Analysis of Concordance (PAC) framework was implemented to assess the translational concordance between preclinical and clinical ASM response for 32 FDA-approved ASMs. Preclinical ASM responses in historically used seizure models were collected. Protective indices based on reported TD_50_ and ED_50_ values were calculated for each ASM in each preclinical model. A weighted scale representing relative anticonvulsant effect was used to grade preclinical ASM response for each seizure model. Data depth was further scored based on the number of evaluated ASMs with publicly available data. Established reports of clinical ASM use in patients with FOS were similarly evaluated and a weighted scale representing prescribing patterns and perceived efficacy used to grade clinical ASM response for each indication. To assess the predictive validity of preclinical models, a unified translational scoring matrix was developed to assign a concordance score spanning the spectrum of complete discordance (-1) to complete concordance (1) between preclinical and clinical ASM responses. Scores were summed and normalized to generate a global translational concordance score.

**Results:** The preclinical models with the highest translational concordance and greatest data depth for FOS were rodent maximal electroshock seizure (MES), mouse audiogenic seizure, mouse 6 Hz (32mA) and rat amygdala kindling.

**Significance:** The PAC-FOS framework highlights mouse MES, mouse audiogenic and mouse 6 Hz (32mA) as three acute seizure models consistently demonstrating high predictive validity for FOS. We provide a pragmatic decision tree approach to support efficient resource utilization for novel ASM discovery for FOS.

**KEY POINT BOX:** Using a newly developed translational scoring matrix, we provide novel insights into the clinical validity of common preclinical seizure models for FOS.

- The PAC-FOS Framework identifies mouse MES, audiogenic and 6-Hz 32 mA as three acute models with greatest predictive validity and versatility for FOS drug discovery.
- We present a pragmatic approach and decision tree to support efficient use of drug discovery resources and in consideration of the 3Rs of animal ethics.
- The work presented would allow for faster and more effective screening of ASMs, while potentially reducing future patient exposures to likely ineffective drugs.

## INTRODUCTION

Epilepsy is a complex neurological disease characterized by unprovoked, spontaneous seizures with a global prevalence of approximately 50 to 60 million people.^1,2^ In the United States, an estimated 3.5 million have an epilepsy diagnosis,^3^ approximately 60% of whom have focal onset seizures (FOS), previously known as partial seizures.^4,5^ FOS refers to seizures originating in networks limited to one hemisphere and that are classified based on patient awareness levels during a seizure. Patients with focal onset impaired awareness seizures (formerly complex partial seizures) typically appear confused or disoriented and may present with unusual, repetitive actions. In focal onset aware seizures (formerly simple partial seizures), awareness is unaffected although patients may be unable to talk or respond. Focal seizures may progress from focal to bilateral tonic-clonic seizures. Temporal lobe epilepsy (TLE) is the most common type of focal epilepsy, beginning in the temporal lobe, and may involve one or both temporal lobes.

Thirty-two antiseizure medications (ASMs) with diverse mechanisms of action are FDA-approved and available in the United States for the treatment of adult epilepsy, including FOS.^6^ ASM efficacy and achievable levels of seizure control tend to vary across seizure types based on the respective mechanisms of action, such that individual ASMs are indicated for specific epilepsy types.^7^ For example, common monotherapies for FOS include the sodium channel blockers, lamotrigine, lacosamide, carbamazepine and oxcarbazepine, as well as levetiracetam, which modulates presynaptic neurotransmitter release by binding to synaptic vesicle protein 2A (SV2A).^8^ Despite the availability of several ASMs, approximately 30-40% of patients are refractory to current treatments,^9,10^ thus highlighting an urgent need for novel treatment options.

Central to the historical discovery of novel ASMs is screening for anticonvulsant activity in preclinical rodent models, predominantly maximal electroshock (MES) and pentylenetetrazole (PTZ) acute seizure models. Several other acute and chronic seizure models have been established and validated for anticonvulsant drug discovery in recent years, including kindling models of acquired TLE and genetic models of spike-wave discharges and absence epilepsy. However, the translational concordance of each model for clinical efficacy across the spectrum of human epilepsies remains unclear. Thus, it is timely to assess the predictive validity of preclinical models to accelerate novel drug discovery efforts.

Optimized resource allocation to streamline drug discovery in a high-throughput manner is a critical translational goal to accelerate early identification of useful and impactful therapies for FOS, while reducing the risk of clinical failure. Furthermore, and given the multitude of potential animal models, there is increasing recognition of the importance of standardization and harmonization of preclinical experimental practices for facilitating appropriate conclusions regarding the therapeutic potential of novel agents.^11^ Here, we sought to establish the translational concordance between ASM response in commonly used preclinical seizure models and focal epilepsy patients to define those animal model(s) with the greatest predictive validity and suitability to early drug discovery campaigns. Concordance findings were evaluated alongside practical considerations including time-intensiveness, resource availability and the 3Rs of animal ethics^12^ – Replacement, Reduction and Refinement – to develop a pragmatic approach, including decision tree, for accelerating drug discovery for FOS.

## METHODS

The Praxis Analysis of Concordance (PAC) framework (**Figure 1**) was implemented to assess the translational concordance between *preclinical ASM response* in commonly used seizure models and *clinical ASM response* in focal epilepsy patients (PAC-FOS) for the 32 FDA-approved ASMs currently available in the United States.^13^

**Figure 1.**
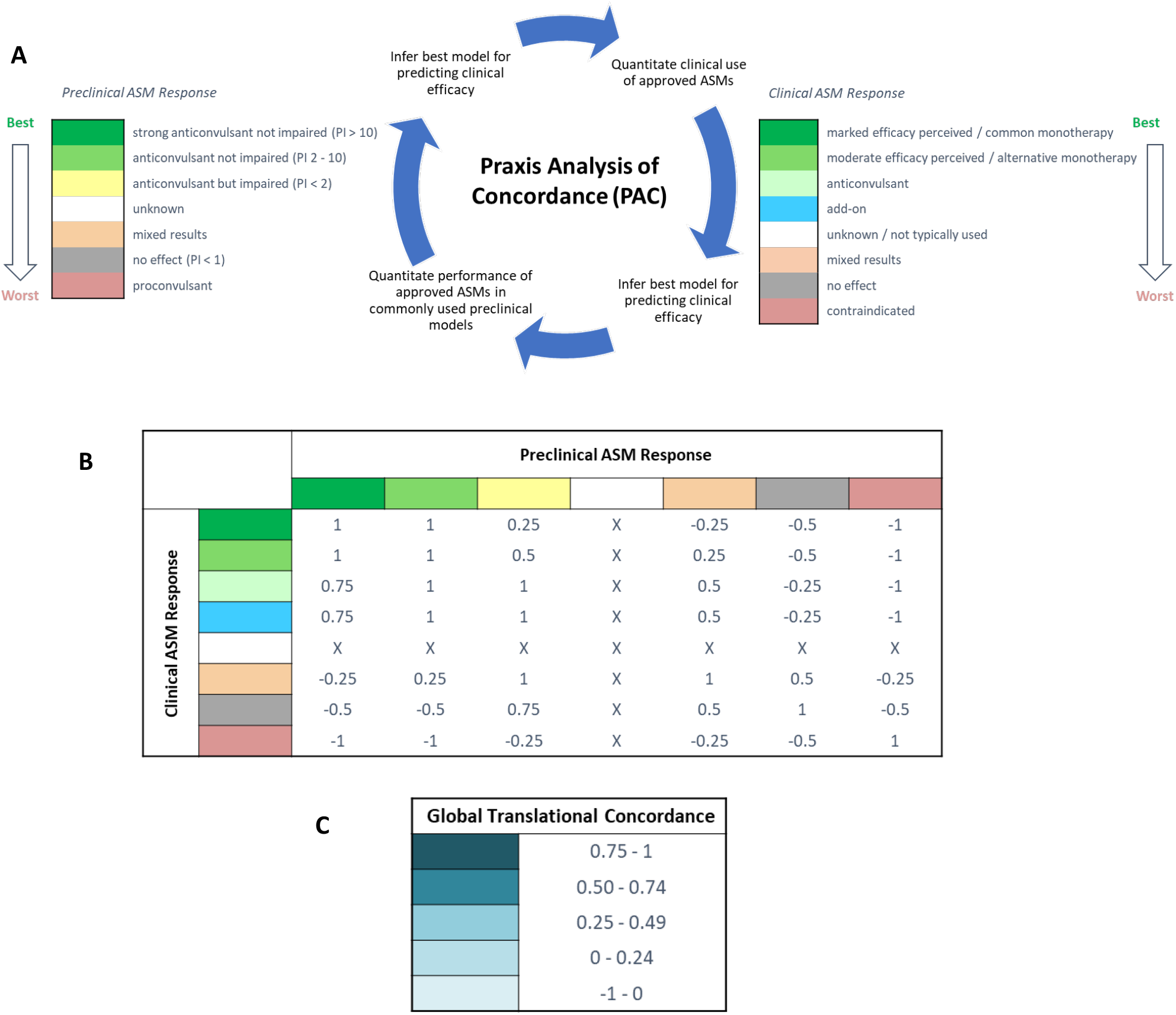
PAC Analysis Framework. (**A**) An overview of the PAC framework developed and applied to assess concordance of ASM response in preclinical models with clinical efficacy in adult patients with focal onset seizures. Performance of approved ASMs in commonly used rodent preclinical seizure models was first evaluated based on reported TD_50_ and ED_50_ values, with *preclinical ASM response* graded for each model according to a weighted scale capturing relative preclinical anticonvulsant effect. Clinical use of approved ASMs was similarly evaluated based on established reports, with *clinical ASM response* graded according to a weighted scale capturing relative clinical anticonvulsant effect. (**B**) A unified scoring matrix was developed to assign translational concordance between *preclinical* and *clinical ASM response*. Values ranged from 1 for complete concordance to -1 for complete discordance. (**C**) For each preclinical seizure model, individual ASM concordance scores were summed and normalized (total translational concordance score/ total number of ASMs with data publicly available) to generate a *global translational concordance score*, weighted from highest (0.75 to 1) to lowest (-1 to 0) concordance. An example application of the PAC framework in FOS is provided in Supplemental Figure 1.

### Preclinical ASM Response

PubMed and the PANAChE database, an NIH National Institute of Neurological Diseases and Stroke (NINDS) resource (http://panache.ninds.nih.gov.), were searched to collect *preclinical ASM responses* in seizure models that have been used historically and/or have been established by the NINDS Epilepsy Therapy Screening Program (ETSP).^14^ Search terms for *preclinical ASM response* included the candidate ASM, seizure model, and species. For each ASM effective in each preclinical seizure model, a protective index (PI) value was calculated from reported median tolerability (TD_50_) and median efficacy (ED_50_) values. Where more than one reference was available, each complete reference was examined, and a representative reference selected to calculate the PI. Studies that reported both efficacy and tolerability values were prioritized in the final analysis. In instances where an ED_50_ and TD_50_ value were not reported in a single study, results from different sources were combined to calculate a PI value, with rotarod TD_50_ values prioritized over those based on motor impairment assays such as open field, Irwin test and minimal motor impairment (MMI). When alternate references were not available, TD_50_ and ED_50_ values from ETSP reports accessed on the NINDS PANAChE database were used. A weighted scale representing relative anticonvulsant effect was used to grade the *preclinical ASM responses* for each seizure model ranging from strong anticonvulsant not impaired (PI > 10) to proconvulsant (**Figure 1A**). Corresponding color-codes were utilized to aid in visualization of graded *preclinical ASM responses*.

### Clinical ASM Response

Searches in PubMed, American Epilepsy Society Annual Meeting abstracts, Epilepsy Foundation and National Institute for Health Care and Excellence websites were performed to collect *clinical ASM responses* for each ASM for FOS that included aware, impaired awareness, focal to bilateral tonic-clonic (secondarily generalized) seizures and TLE. Published reports of ASM use and perceived efficacy and prescribing patterns were evaluated in adult patients with focal epilepsies, and a weighted scale representing relative anticonvulsant effect was used to grade *clinical ASM responses* ranging from marked efficacy perceived and/or common monotherapy to contraindicated (**Figure 1A**). Corresponding color-codes were utilized to aid in visualization of graded *clinical ASM responses*.

### Translational Concordance Scoring

To assess the predictive validity of preclinical seizure models across individual ASMs, a unified scoring matrix was developed (**Figure 1B**). Using this matrix, a *translational concordance score*, capturing the spectrum of complete discordance (-1) to complete concordance (1) between *preclinical* and *clinical ASM responses*, was assigned for each preclinical model and clinical combination. Assigned scores were then summed and normalized to generate a *global translational concordance score* (**Figure 1C**). Given the variability in data available for collation of *preclinical ASM responses*, and to aid interpretation of findings, data depth was similarly scored on a weighted scale from robust to minimal, based on the number of ASMs tested in each preclinical model, and for which data were publicly available for collation and analysis.

## RESULTS

### PRECLINICAL ASM RESPONSE

The preclinical efficacy of each of the 32 FDA-approved ASMs was evaluated across 23 seizure models: mouse and rat MES, mouse audiogenic, mouse and rat subcutaneous PTZ (scPTZ), zebrafish PTZ, mouse bicuculline and picrotoxin, mouse and rat flurothyl, rat pilocarpine preventative and interventional, GAERS or WAG/Rij, mouse and rat 6 Hz (32 and 44 mA), rat systemic kainate (limbic seizures), mouse intra-hippocampal and intra-amygdala kainate (IHK and IAK), 60 Hz corneal-kindled mouse, and rat hippocampal, amygdala and lamotrigine (LTG)-resistant amygdala kindling (**Figure 2, Supplemental Figure 2**). Based on PI values for each ASM, *preclinical ASM response* for each preclinical seizure model was graded ranging from proconvulsant (red) to strong anticonvulsant not impaired (dark green, PI > 10). ASMs with mixed results and those with no data available were graded accordingly (orange and white, respectively). Not all ASMs tested in the zebrafish PTZ model had corresponding tolerability values. In those cases, *preclinical ASM responses* were graded according to the following scale: ED_50_ < 20 μM (dark green; strong anticonvulsant not impaired), ED_50_ 25-250 μM (light green; anticonvulsant), ED_50_ > 250 μM (yellow; anticonvulsant but impaired) and ED_50_ > 10 mM (gray; no effect).

**Figure 2.**
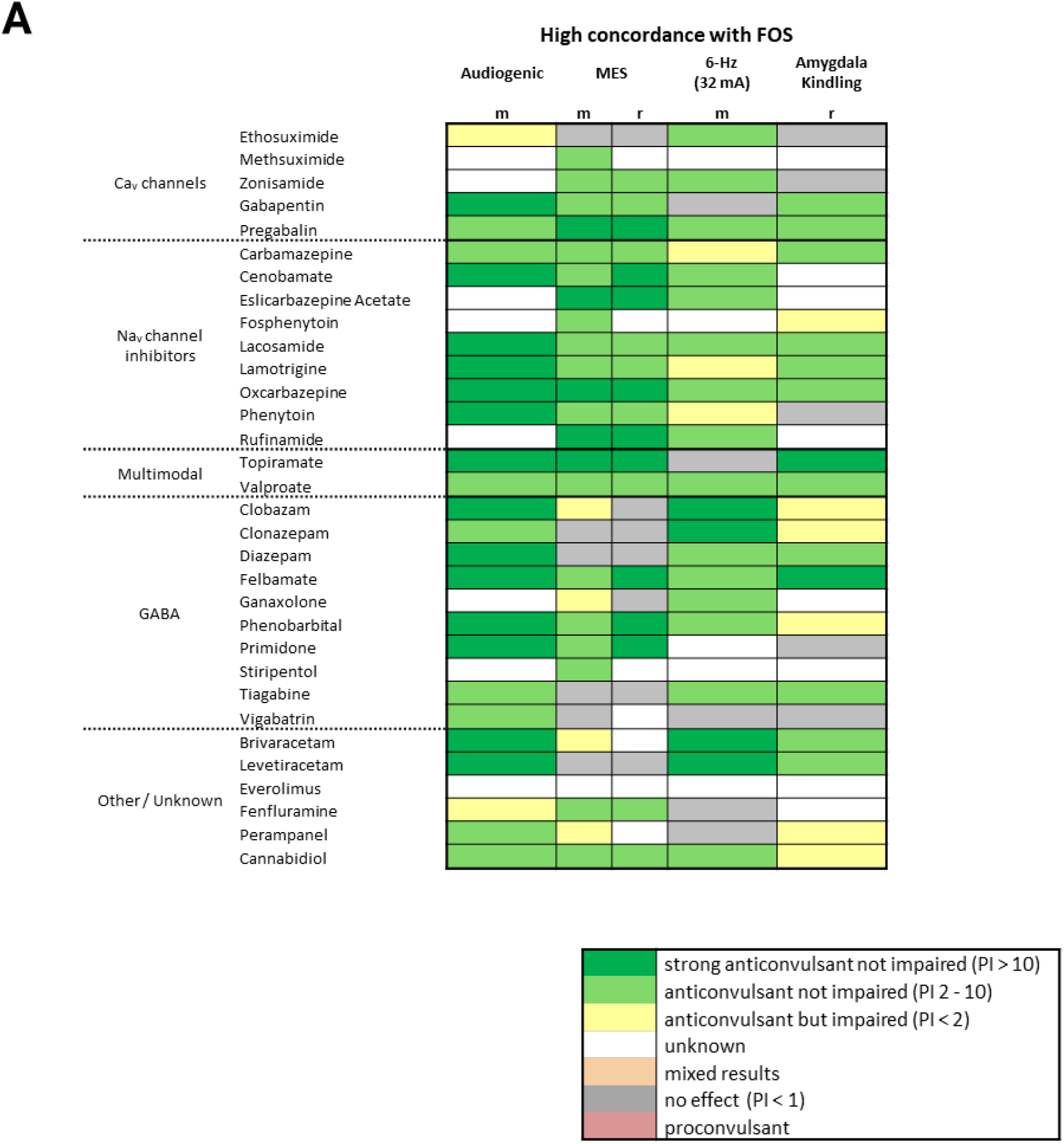
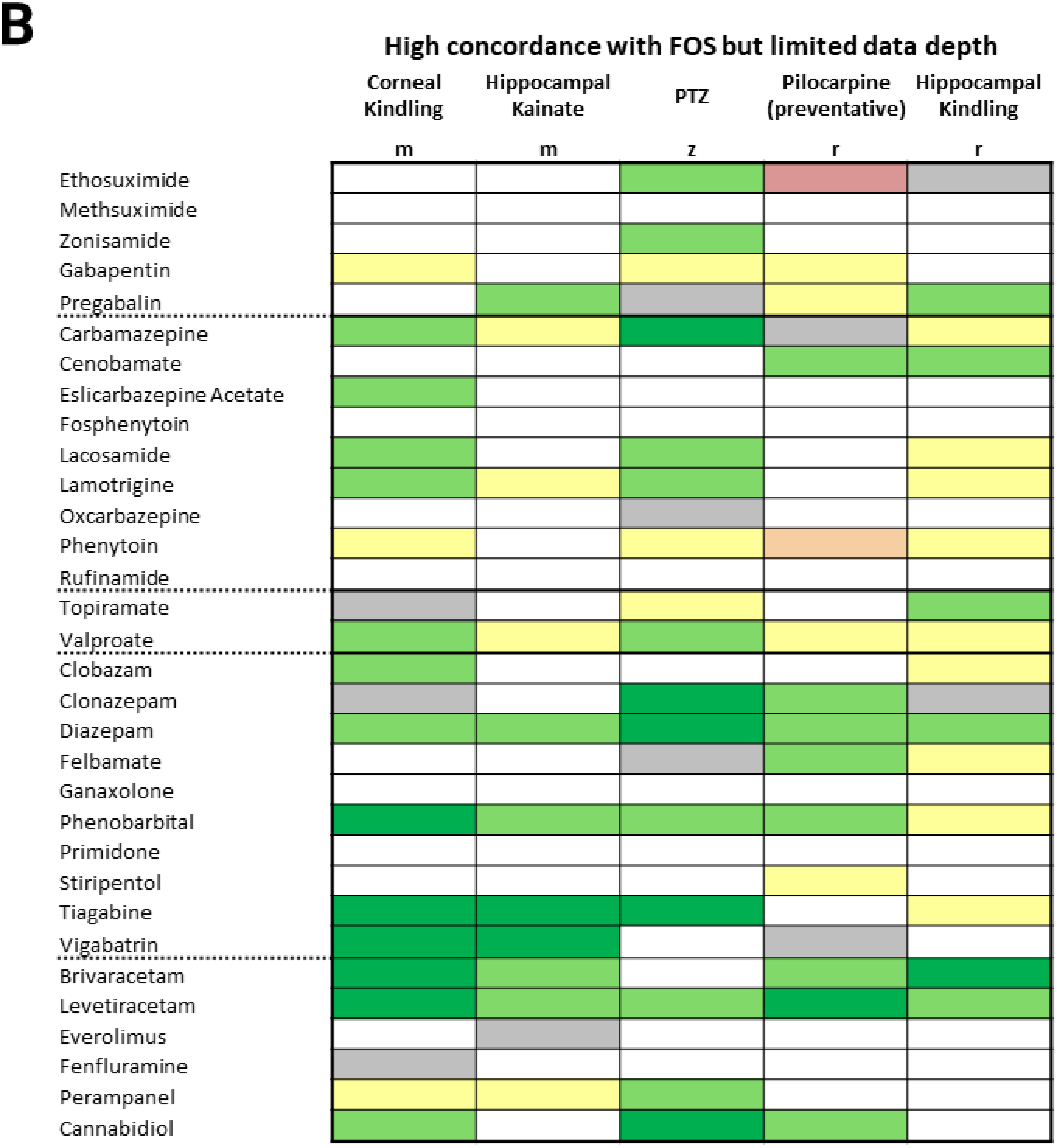
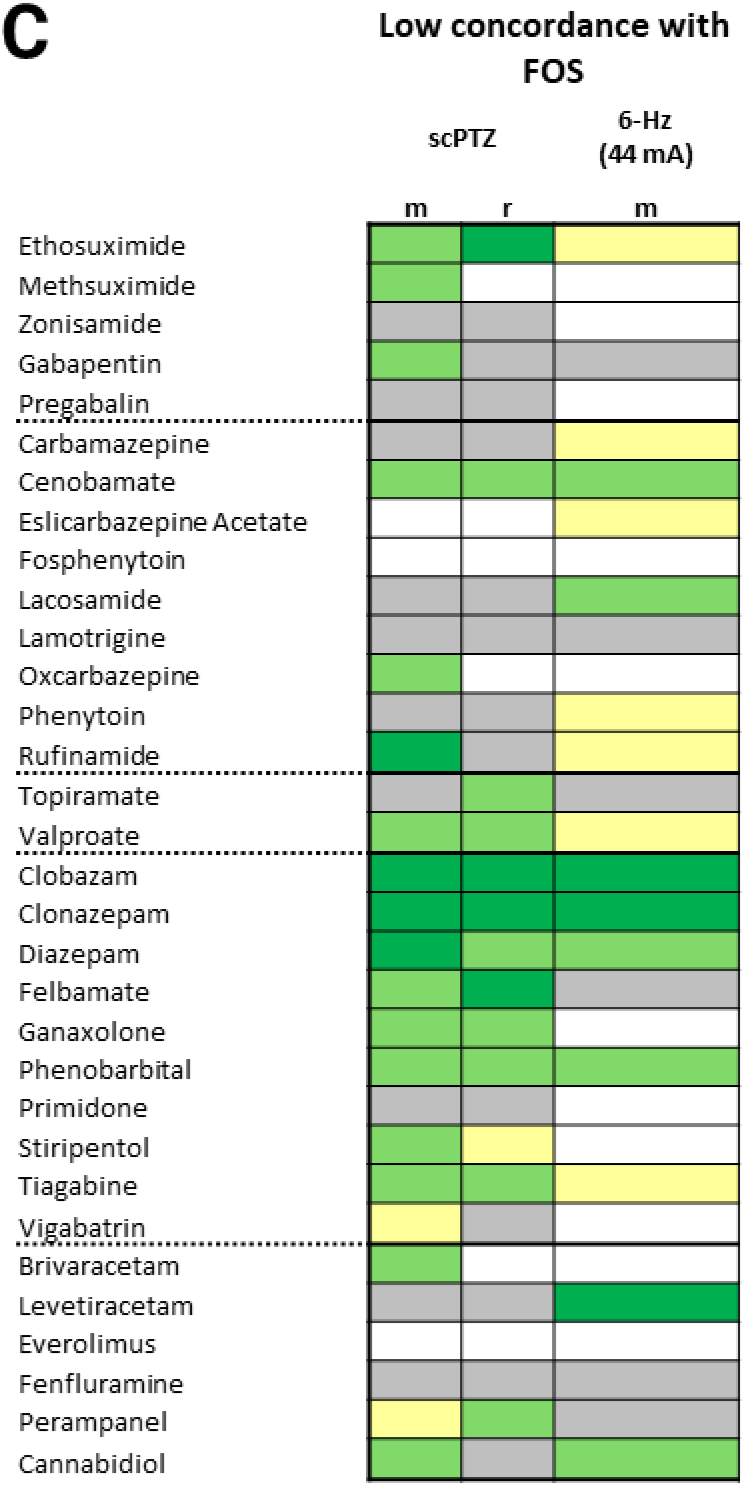
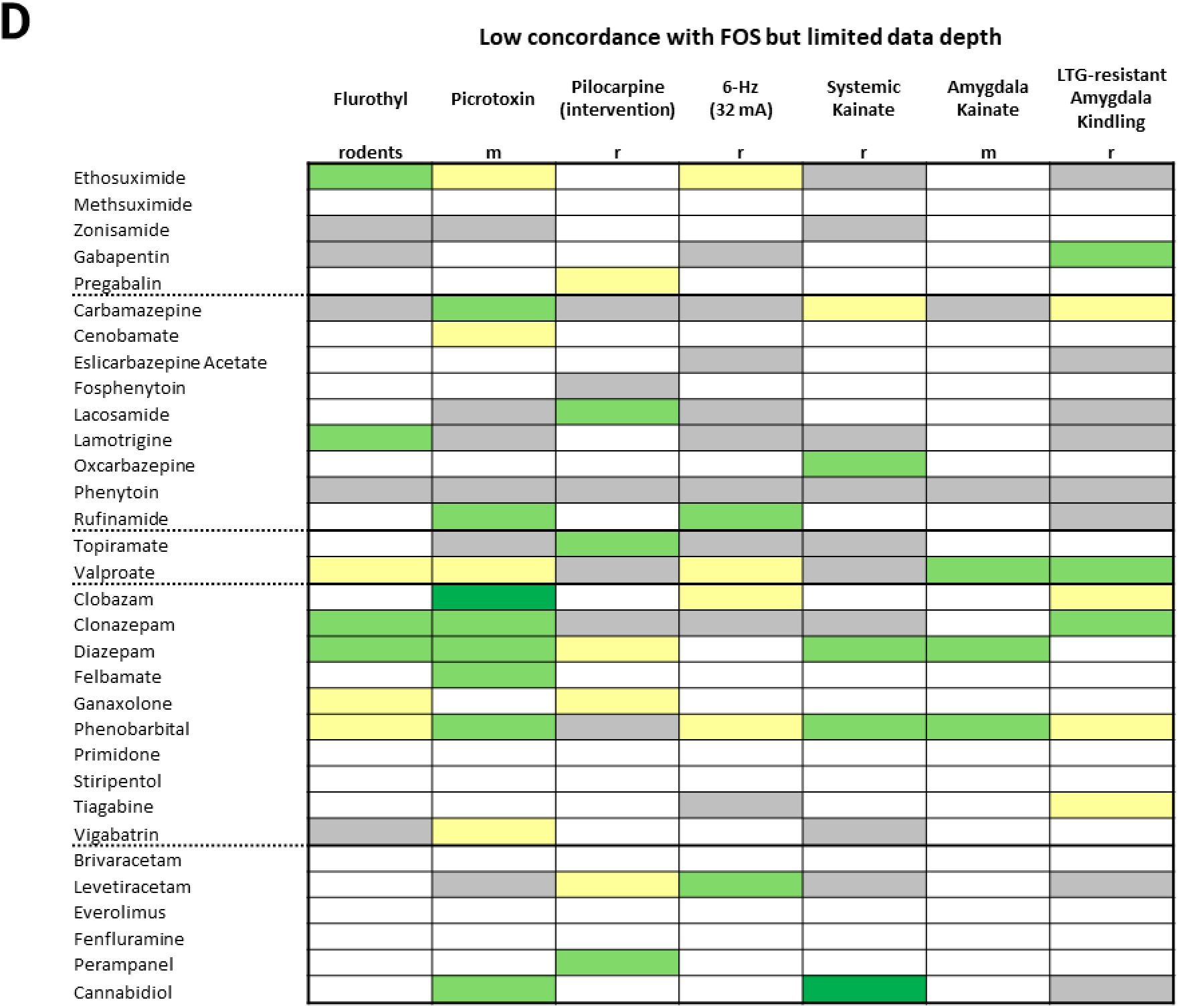
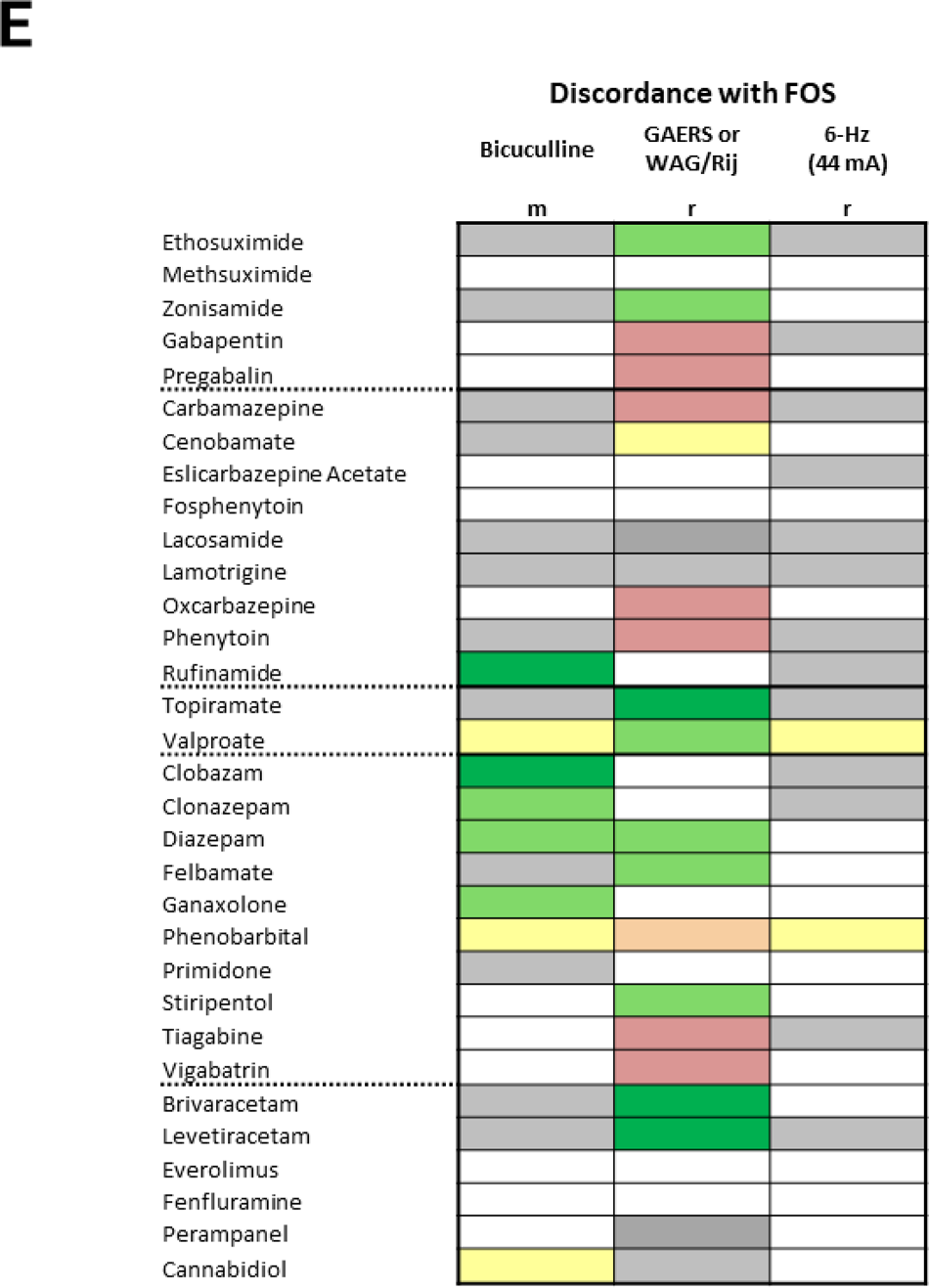
Preclinical ASM Response. The preclinical efficacy of 32 FDA-approved ASMs currently available in the US was examined in a total of 23 seizure models across multiple species. Colors denote grading of *preclinical ASM response* based on reported TD_50_ and ED_50_ values for each model to calculate a protective index (PI), resulting in a weighted scale capturing relative preclinical anticonvulsant potential (*inset key shown in* **A**). Data depth was similarly scored on a weighted scale from most robust to minimal, based on the number of approved ASMs that have been tested with data publicly available for collation in each model. Preclinical seizure models were grouped according to class/mechanism of action: calcium and sodium channel blockers, multimodal agents, GABAergic agents as well as agents with other mechanisms of action (including mTOR inhibitors, modulators of SV2A, selective serotonin reuptake inhibitors, and AMPA inhibitors) and those exhibiting **A**) high concordance with FOS, **B**) high concordance with FOS but with limited data depth, **C**) low concordance with FOS, **D**) low concordance with FOS but with limited data depth, and **E)** discordance with FOS. Please refer to Supplemental Figure 2 for individual data sources evaluated to assign depicted gradings. m, mouse; r, rat; z, zebrafish; MES, maximal electroshock seizure, PTZ, pentylenetetrazole; preventative, treatment administered before pilocarpine-induced *status epilepticus*; interventional, treatment administered after pilocarpine-induced *status epilepticus*.

### CLINICAL ASM RESPONSE

Perceived clinical efficacy and prescribing patterns were evaluated for each of the 32 FDA-approved ASMs across the following established ILAE focal epilepsy classifications:^15^ focal onset aware, focal onset impaired awareness, focal to bilateral tonic-clonic (secondarily generalized) seizures and temporal lobe epilepsy (**Figure 3, Supplemental Figure 3**). Based on published reports, *clinical ASM response* for each clinical indication was graded to capture the relative clinical anticonvulsant potential for each ASM, ranging from contraindicated (red) to marked efficacy perceived / common monotherapy (dark green).

**Figure 3.**
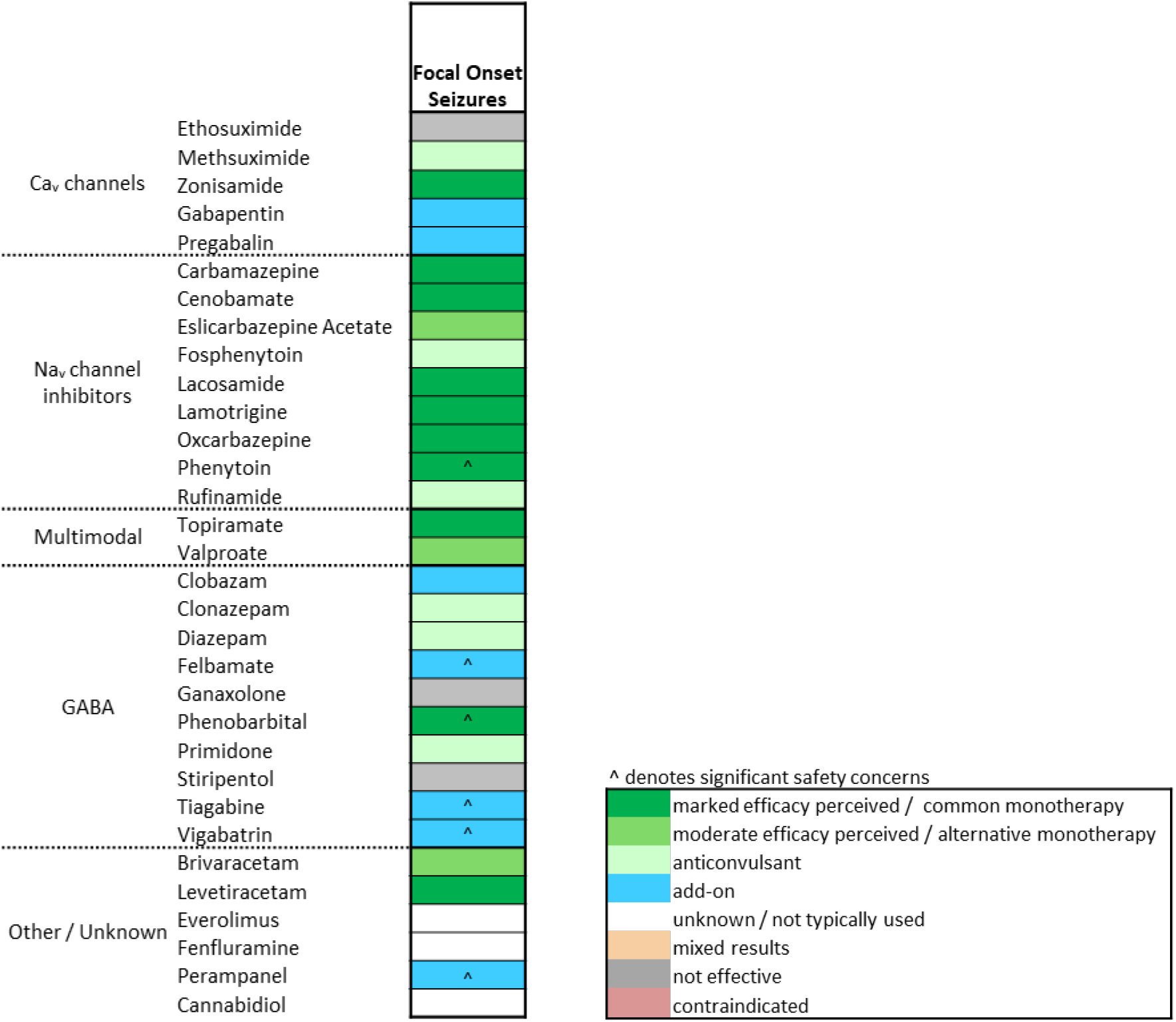
Clinical ASM Response. Clinical efficacy of the same 32 FDA-approved ASMs was evaluated based on established reports of perceived efficacy and use. Colors denote grading of clinical ASM response based on prescribing patterns for FOS, resulting in a weighted scale capturing relative clinical anticonvulsant potential (*inset key*). Please refer to Supplemental Figure 3 for individual data sources evaluated to assign depicted gradings.

### SUMMARY OF TRANSLATIONAL CONCORDANCE FINDINGS

Global translational concordance results and corresponding data depth are summarized in **Figure 4**. The audiogenic seizure model had the highest concordance with FOS, with a global translational concordance score of 0.9. Additional preclinical models with high (> 0.5) global translational concordance were MES (mouse and rat), zebrafish PTZ, preventative pilocarpine, mouse 6 Hz (32 mA), hippocampal kainate, 60 Hz corneal kindling, and hippocampal and amygdala kindling. However, many of these preclinical models have limited data depth engendering uncertain confidence in their global translational concordance. Models where less than two-thirds of approved ASMs have been tested with data publicly available for collation were excluded to restrict analysis to models with robust data-depth. This highlighted four models with robust data depth and high concordance with FOS: audiogenic, MES, mouse 6 Hz (32 mA), and rat amygdala kindling (**Figure 2A**).

**Figure 4.**
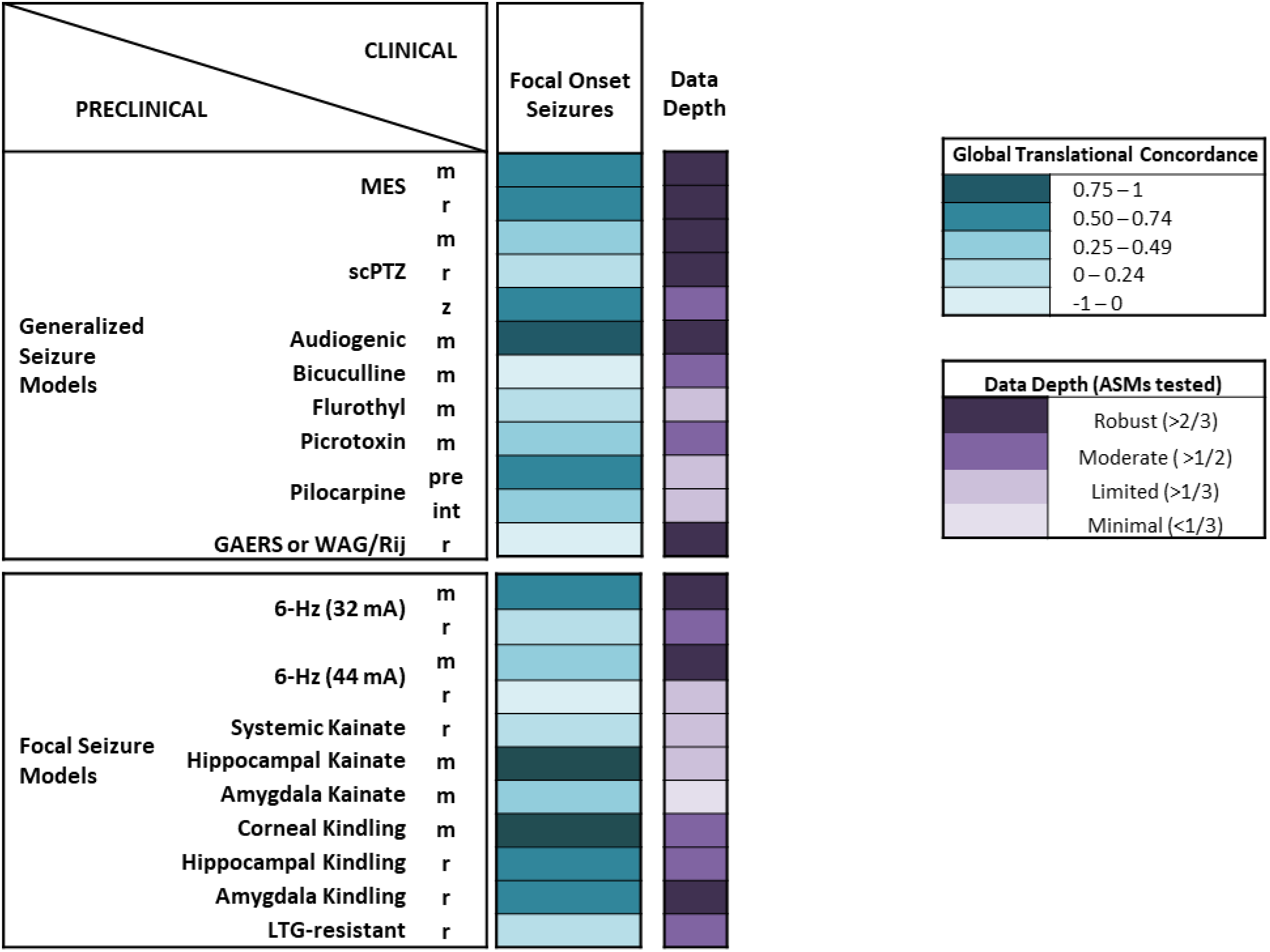
FOS Global Translational Concordance. Global translational concordance of each preclinical seizure model for FOS and associated data depth. Teal shading corresponds to a weighted scale from highest (0.75 to 1) to lowest (-1 to 0) concordance scores. Data depth (purple shading) was similarly scored on a weighted scale from most robust to minimal, based on the number of approved ASMs that have been tested with data publicly available for collation in each model. Data depth shading thus denotes the proportion of all 32 approved ASMs, from robust (more than two-thirds of all 32 ASMs having data publicly available for analysis) to minimal (less than one-third of all 32 ASM having data publicly available for analysis).

#### Models with high concordance

Audiogenic-induced seizures in Frings or DBA/2J inbred mouse strains exhibited high concordance with FOS. The first observations of audiogenic-induced seizures were made by Pavlov in St. Petersburg in the early 1920s and in the Wistar Institute in Philadelphia. Two inbred mouse strains, Frings and DBA/2J, exhibit a susceptibility to audiogenic-induced seizures^16–18^ Exposure to a high-intensity sound stimulation as either a pure 8-16 kHz tone, a broad-band noise (e.g. a bell ring) or white noise will trigger seizures characterized by wild running, loss of righting reflex and tonic hindlimb extension.^19^ In both mouse strains, seizure susceptibility becomes apparent around weaning. While Frings mice remain susceptible to audiogenic-induced seizures throughout their life, the window of susceptibility for DBA/2J mice is narrow (∼P18-30) due to progressive hearing loss and deterioration to total deafness. Treatments are administered prior to seizure induction and are evaluated for the ability to protect from tonic hindlimb extension. In this study, all classes of ASMs showed efficacy in an audiogenic seizure model; albeit with a broad range of PI values, from < 2 to upwards of 40 (**Figure 2A**).

While MES-induced seizures are electrophysiologically consistent with generalized tonic-clonic seizures, MES in both mice and rats showed high concordance with FOS. MES is the most widely used preclinical seizure model, is highly reproducible across labs and is one of few models that have been clinically validated.^20–23^ As the name suggests, the MES test applies a short, high frequency electrical current through transcorneal or transauricular electrodes to induce acute seizures characterized by tonic extension of forelimbs and hindlimbs followed by brief episodes of clonic activity. MES was sensitive to several mechanistic classes, with sodium channel blockers tending to perform best (**Figure 2A**). Compounds that modulate GABAergic signalling typically demonstrated lower effectiveness, exhibiting no effect or only anticonvulsant efficacy at motor impairing doses. Notably, levetiracetam is ineffective in the MES acute seizure model.^24^

Our findings demonstrate similarly high concordance between the mouse 6 Hz (32 mA) acute seizure model and FOS. The 6 Hz acute seizure model was introduced in the 1950s as a focal seizure model,^25,26^ but its development was originally abandoned because several clinically useful drugs were ineffective, calling into question its clinical relevance. Decades later, the model was revisited and has become a widely implemented acute seizure model for screening compounds.^27,28^ A low frequency (6 Hz), long duration (3 s), 32 mA-equivalent (1.5x CC_97_) transcorneal or transauricular stimulation is applied to induce psychomotor seizures characterized by immobility, jaw clonus, forelimb clonus, stereotyped movements, twitching of the vibrissae and/or Straub tail. Several classes of ASMs have anticonvulsant efficacy in the mouse 6 Hz acute seizure mouse model at 32 mA (**Figure 2A**). In 6 Hz, GABAergics show strong anticonvulsant effect without impairment of function while sodium channel blockers are less efficacious or showefficacy only at motor impairing doses.

Lastly, the amygdala kindling model also exhibited high concordance with FOS. In the amygdala kindling model, an electrical stimulation (50 Hz, 2 s train of 1 ms biphasic 200 μA pulses) is repeatedly administered until seizure criterion of at least 5 consecutive fully kindled clonic seizures with rearing and/or falling, Racine Stage 4 or 5 is reached.^29,30^ Sodium channel blockers and GABAergics tend to suppress motor-seizure severity at doses devoid of behavioral impairment (**Figure 2A**). Furthermore, third generation ASMs including levetiracetam, brivaracetam and felbamate demonstrate anticonvulsant efficacy in the amygdala kindling model.

#### Models with high concordance but limited data-depth

Electrical hippocampal and amygdala kindling models were adapted in the early 1980s to apply electrical stimulation to the corneas of mice. Our PAC-FOS findings revealed high concordance between the mouse corneal kindling (60 Hz) model and FOS, however, only 59% (19/32) of approved ASMs have data publicly available for this model. Electrical hippocampal and amygdala kindling models were adapted in the early 1980s to apply electrical stimulation to the corneas of mice. Notably, adaptations of the 60 Hz corneal stimulation paradigm have been established in recent years to evoke pharmacoresistant seizure models using variations in the electrical current or frequency applied,^31–33^ which may be well-suited to identify treatments for drug-resistant FOS. Regardless of stimulation paradigm, the mouse corneal kindling model is evoked with a stimulation protocol delivered twice daily for 2-4 weeks until at least 5 consecutive fully-kindled seizures (clonic seizures with rearing and loss of posture, Racine stage 5) are consistently elicited.^34^ Treatments are administered to fully kindled mice prior to the kindled stimulation, and therapeutic efficacy is assessed by the ability to protect from generalized motor seizures. Most ASMs show anticonvulsant activity against generalized motor seizures in the corneal kindling model, with a large range of PI values (**Figure 2B**).

The intrahippocampal kainate (IHK) mouse model exhibited high concordance with FOS, however, less than half of all approved ASMs have data publicly available for this model. The IHK mouse model of mesial temporal lobe epilepsy (MTLE) is a validated model that recapitulates many histological, electrophysiological, behavioral and pharmacological features of TLE patients.^35–37^ In the IHK mouse model, a non-convulsive *status epilepticus (SE)* lasting several hours is induced. Epileptogenesis then occurs over the 2-3 weeks following this initial insult and mice develop spontaneous electrographic hippocampal paroxysmal discharges (HPDs) that are associated with behavioral arrest and/or mild motor automatisms. The HPDs are defined as rhythmic high-amplitude sharp waves at 5-10 Hz lasting 5-20 seconds that occur approximately 30-60 times per hour.^35–37^ As our findings indicate (**Figure 2B**), sodium channel blockers, such as carbamazepine and lamotrigine, do reduce HPDs, but only at doses that are motor impairing. In contrast, GABA mimetics and the third generation ASMs levetiracetam, brivaracetam and perampanel suppress HPDs at doses devoid of behavioral impairment.

The zebrafish PTZ model demonstrated high concordance with FOS, but only 59% (19/32) of approved ASMs have been reportedly tested, with tolerability data not available for many ASMs (**Figure 2B**). The GABA_A_ receptor antagonist PTZ (Metrazol) is the most widely used chemoconvulsant in rodents as a model of generalized clonus. More recently, the PTZ test has been adapted for zebrafish to generate a higher throughput model,^38^ wherein PTZ is added directly to the aquaculture media of zebrafish larvae. The resulting seizures are characterized by rapid whirlpool-like movements and rapid jerky swimming behavior with bouts of body-stiffening and loss of posture.^39^ The anticonvulsant efficacy of several ASM classes is evident, however, there is variability in the effects of ASMs in the zebrafish PTZ model compared with mouse and rat subcutaneous PTZ (scPTZ) acute seizure models (**Figure 2B & 2C**). This variability may be a result of scarce zebrafish tolerability datasets, or it may reflect true species variability that could be a useful differentiator for screening cascades.

There was high concordance between FOS and the preventative pilocarpine *SE* model, although data depth was limited with only half of ASMs having data publicly available. In this model, Sprague-Dawley rats are first injected with lithium chloride followed by the muscarinic acetylcholine receptor agonist pilocarpine to induce *SE*.^40^ Our PAC findings demonstrate the large variability in the efficacy of ASMs to prevent pilocarpine-induced *SE* (**Figure 2B**). Of note, compounds that modulate GABAergic signalling tend to effectively prevent pilocarpine-induced *SE*. On the other end of the spectrum, ethosuximide and acetazolamide are proconvulsant in this model, tending to exacerbate *SE*.

There was also high concordance between the hippocampal kindling model and FOS, but data depth was again limited, with only 53% (17/32) of ASMs tested. Similar to the amygdala kindling model, the hippocampal kindling model repeatedly administers an electrical stimulation until rats are fully-kindled. Collated *preclinical ASM responses* highlighted cenobamate and diazepam as efficacious in the hippocampal kindling model, but most sodium channel blockers and GABA mimetics only show efficacy at motor impairing doses (**Figure 2B**). Contrastingly, the third generation ASMs levetiracetam, brivaracetam, pregabalin and topiramate suppress secondarily generalized motor seizures at doses devoid of behavioral impairment.

#### Models with low concordance

Ten preclinical seizure models demonstrated low (<0.5) translational concordance scores for FOS, seven of which had limited data depth (**Figure 2C and D**): scPTZ (mouse and rat), mouse 6 Hz (44 mA), flurothyl, picrotoxin, pilocarpine *SE* (interventional), systemic kainate, rat 6 Hz (32 mA), amygdala kainate (IAK), and LTG-resistant amygdala kindling.

#### Discordant models

Interestingly, there was discordance between FOS and bicuculline and rat 6 Hz (44 mA) models (**Figure 2D**). Unsurprisingly, discordance was observed between FOS and genetic models of absence epilepsy (Genetic Absence Epilepsy Rats from Strasbourg (GAERS) or inbred Wistar Albino Glaxo Rats from Rijswijk (WAG/Rij). Common FOS therapies were not effective in bicuculline and rat 6 Hz (44 mA) acute seizure models, while several ASMs including carbamazepine, phenytoin and gabapentin demonstrated proconvulsant effects in the Genetic GAERS and/or WAG/Rij models.^41–43^

## DISCUSSION

To the best of our knowledge, this study represents the first comprehensive examination of the translational concordance between commonly used preclinical animal models across the clinical epilepsy spectrum. Following evaluation of published reports of *preclinical* and *clinical ASM responses*, we applied the PAC framework employing a novel translational scoring matrix gated by PI thresholds to identify key preclinical models with the highest predictive validity for FOS (PAC-FOS).

Focusing only on models with robust data depth, a clear pattern of concordance was revealed. Preclinical models with the highest translational concordance for FOS were found to be audiogenic seizures, MES (mouse and rat), mouse 6 Hz (32 mA) and rat amygdala kindling. MES findings were consistent across mouse and rat models, suggesting one model is likely sufficient for early-stage drug discovery. Therefore, the mouse model was selected in the decision tree as it is a more efficient, cost-effective option involving less investigational compound and simpler husbandry requirements.

Based on our findings, we present a decision tree for FOS drug discovery that supports efficient use of resources and takes into consideration the 3Rs of animal ethics, wherein an optimized workflow (**Figure 5**) involves the following three acute seizure models: audiogenic, mouse MES and mouse 6 Hz (32 mA). We propose that progression to development should be considered for compounds with minimum PI values > 2 in two of these models. Notably, all 13 ASMs with moderate to marked efficacy perceived in FOS are captured with such a screening cascade, including levetiracetam which is routinely cited as an example of an anticonvulsant “missed” by MES.

**Figure 5.**
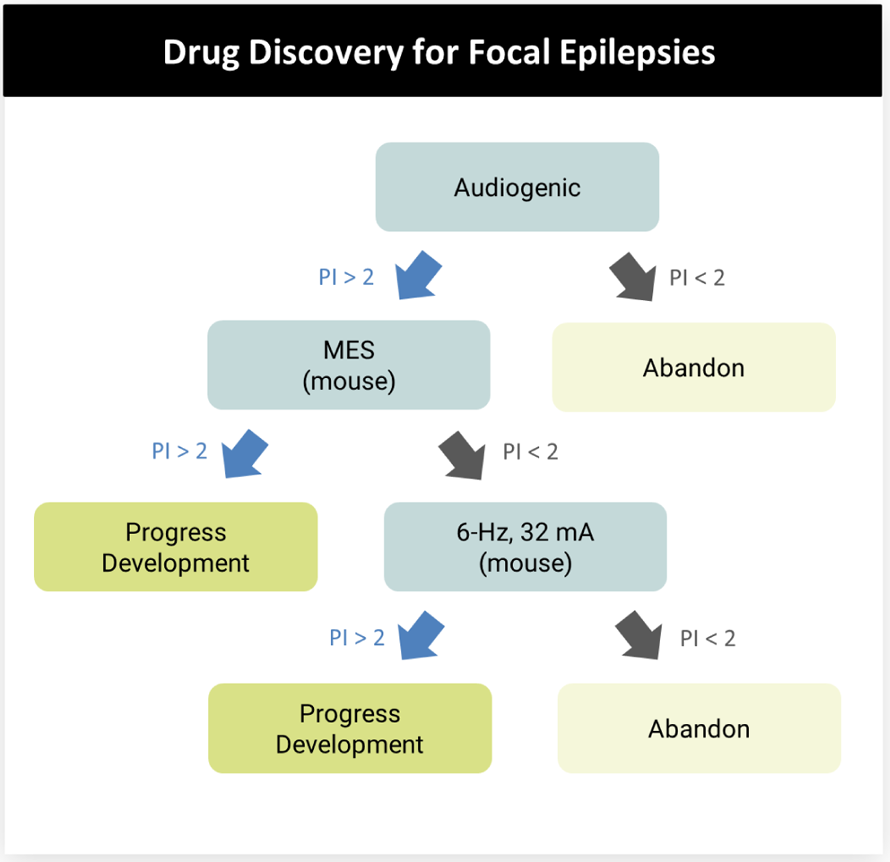
FOS Drug Discovery Decision Tree. PI=protective index value (TD_50_/ED_50_)

The ETSP regards Frings audiogenic seizure models as legacy models, primarily useful in proof-of-concept and screening studies, given Frings mice exhibit a broad response profile to approved ASMs. Indeed, our findings reinforce the notion that audiogenic seizure susceptible models alone are unlikely to be useful for identifying false positives and determining clinical positioning. For instance, ethosuximide and phenytoin are active against sound-induced seizures despite markedly different clinical utility across the epilepsy spectrum. Addressing this issue, the PAC framework and associated PAC-FOS decision tree ensures progression beyond the audiogenic seizure model is gated to PI thresholds, thereby enabling compounds such as ethosuximide (PI = 1) to be distinguished from those such as phenytoin (PI = 10.5). Following the proposed cascade, the broad response profile for compounds in audiogenic seizure models essentially casts a large net (reducing false negatives) for compounds with anticonvulsant potential for FOS. PI thresholds rather than binary responses can then be used to reduce false positives and identify compounds with the greatest chance of successful progression along the drug discovery pipeline. Considerations surrounding the relative efficiency of the audiogenic seizure model relate to the narrow age of use for DBA/2J mice (∼P18-30) versus the potential for mice to be reused two to three times in MES and 6 Hz models without impacting seizure response. Additionally, while Frings mice remain susceptible to audiogenic-induced seizures throughout the lifespan, their access is limited through the University of Utah, further impacting feasibility of the audiogenic model as an exclusive first step in a screening cascade.

The rat amygdala kindling model was also identified as exhibiting high translational concordance with FOS and provides a direct focal seizure assessment as measured by the after-discharge duration and after-discharge threshold. However, the amygdala kindling model is time consuming, costly and laborious, often requiring advanced surgical and EEG techniques. The experimental animals are also required to be housed individually, which makes the model resource intensive to early screening. Our findings suggest that the translational concordances of the audiogenic, MES and 6 Hz models are equivalent to the translational concordance of the amygdala kindling model, such that the proposed drug discovery workflow favors the former acute models as more efficient, cost-effective options.

Findings from the PAC analysis were limited by the depth of publicly available data. Only three agents (carbamazepine, phenobarbital and valproate), that is less than 10% of all approved ASMs, have been tested in all 23 preclinical models. Further, less than half of preclinical models passed the data robustness threshold requiring testing with at least 2/3 of approved ASMs. As such, models with high concordance, yet low data depth should not yet be prioritized in early discovery campaigns, but may provide important insights into lead compounds and potential drug candidates. One notable example, the corneal kindling model, exhibited high translational concordance but had limited data depth. As additional data becomes available, we anticipate the FOS drug discovery decision tree to be further refined and/or additional branches added. In such instances, if the observed high translational concordance of the corneal kindling model holds with a robust dataset, one could envision substituting it for 6 Hz or adding it to the bottom of the decision tree. The same might be applicable for models in the “differentiation phase” of the ETSP workflow. Additional future refinement of the presented decision tree may necessitate reevaluation and adjustment of PI thresholds to prioritize advancement of agents with truly novel therapeutic potential over “me-too” compounds.

It is worth noting that no single preclinical model is homologous in its ability to replicate every aspect of the human disorder, with existing models demonstrating variable resemblance to human FOS. Indeed, MES and audiogenic seizure models demonstrate high translational concordance with FOS despite inducing generalized seizures. As such, our PAC assessment was agnostic to the seizure type induced by the model, with preclinical predictive validity findings for FOS found to be independent of seizure type induced by the model. Parallel efforts are ongoing to evaluate the concordance between commonly used preclinical models and generalized epilepsies, thus extending applicability of the PAC framework to the spectrum of human epilepsies.

### Limitations and Future Directions

One potential limitation associated with this analysis relates to the heterogeneity across preclinical studies including, but not restricted to, varying rodent strains, sexes, dose formulations and pretreatment times. For example, 6 Hz studies have demonstrated high variability with efficacy greatly impacted by mouse strain and experimental conditions as evidenced by a side-by-side comparison of valproic acid, phenobarbital and levetiracetam across two mouse strains within the same laboratory.^31^ Where possible, collation of *preclinical ASM responses* prioritized studies with consistent experimental conditions; however, heterogeneity in mouse strain, sex and age inclusion criteria was an inherent limitation for animal models where publicly available data were scarce. Another limitation involves the determination of *preclinical ASM response* based on published dose (TD_50_ and ED_50_) rather than using measured exposures, which may not be dose-linear. Despite these limitations, our drug discovery workflow for FOS presents several strengths. The PAC framework and associated decision tree was optimized for high specificity and, promisingly, demonstrates both high specificity (1.0; false positive rate 0.00) and high sensitivity (0.92). Perampanel and vigabatrin were the only false negatives, both of which exhibit efficacy for FOS as add-on treatments but are also accompanied by safety issues. Importantly, this study provides a newly developed scoring matrix to assess translational concordance and predictability to provide novel insights into the clinical validity of commonly used preclinical seizure models for FOS. We anticipate these findings to have significant implications for supporting ongoing research and development efforts as well as promoting efficient resourcing for novel ASM development for FOS.

## ACKNOWLEDGEMENTS

The authors acknowledge Sarah Hulls for medical writing and editorial contributions provided in accordance with Good Publication Practice (GPP3).

