## Supplemental Figure 1 for "PAC – A novel translational concordance framework identifies preclinical seizure models with highest predictive validity for clinical focal onset seizures"

**Supplemental Figure 1. Example Application of PAC Framework.** The preclinical efficacy of 32 FDA-approved ASMs currently available in the US was examined in a total of 23 seizure models across multiple species. In this example, grading of *preclinical* (MES mouse model) to *clinical ASM response* is depicted*.* *Preclinical ASM response* was based on reported TD_50_ and ED_50_ to calculate a protective index (PI), resulting in a weighted scale capturing relative preclinical anticonvulsant potential. *Clinical ASM response* was evaluated based on established reports of perceived efficacy and prescribing patterns for FOS, resulting in a weighted scale capturing relative clinical anticonvulsant potential. A unified scoring matrix was developed to assign translational concordance between *preclinical* and *clinical ASM response* for each ASM*.* Values ranged from 1 for complete concordance to -1 for complete discordance. Individual ASM concordance scores were summed and normalized (total translational concordance score/ total number of ASMs with data publicly available) to generate a *global translational concordance score*, weighted from highest (0.75 to 1) to lowest (-1 to 0) concordance. In this example, the total translational score of 19.75 was normalized according to 29 ASMs with data available in the mouse MES model, yielding a global translational concordance value of 0.68. ASM, antiseizure medication; FOS, focal onset seizures; m, mouse; MES, maximal electroshock seizure.

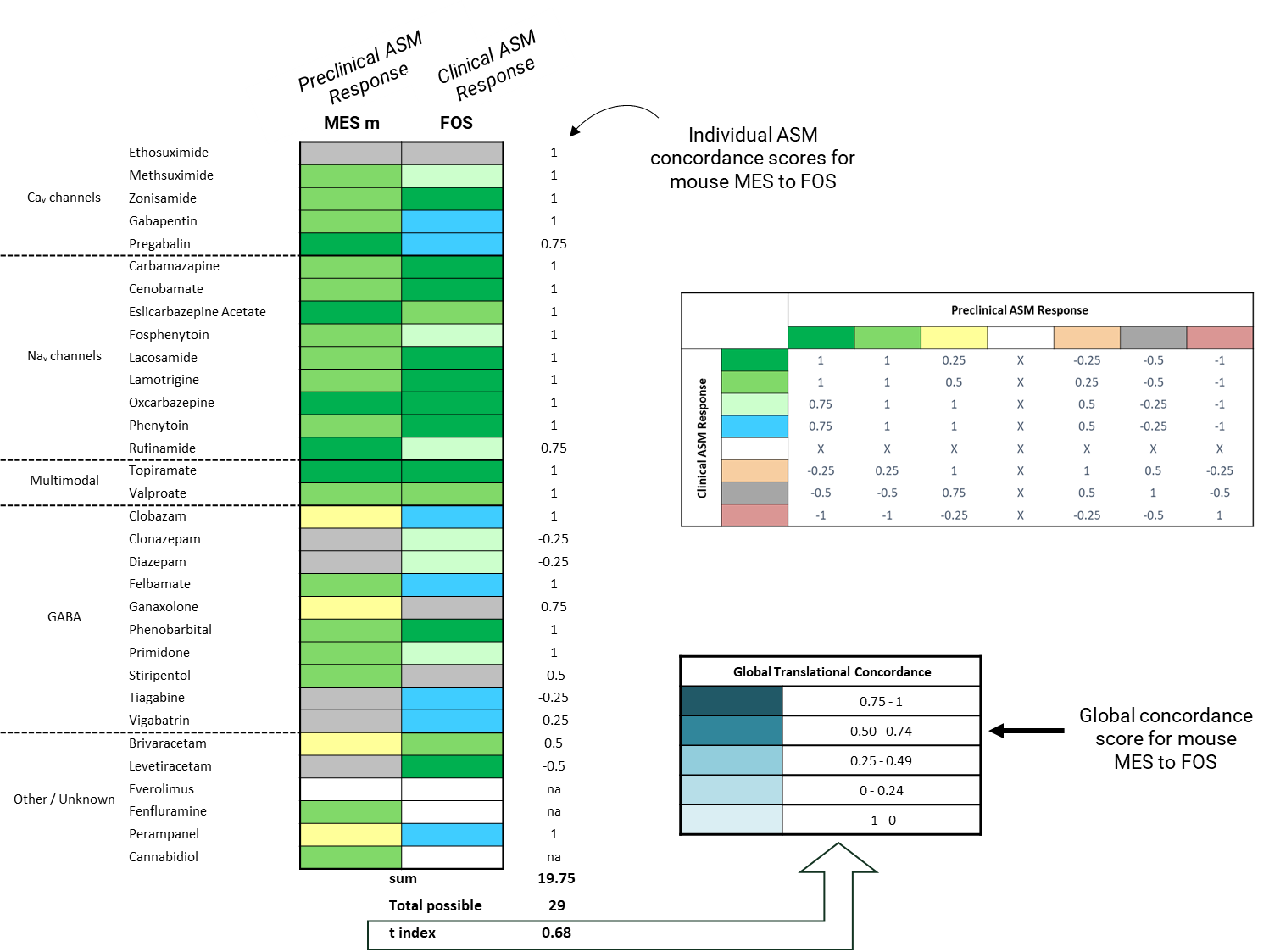

6. Program, A. S. Ethosuximide Test 16 Results - Anticonvulsant Quantification (Frings Mice) (12) Anticonvulsant Screening Program.

7. Program, A. S. Ethosuximide Test 4 Results - Mice I.P. Quantification Anticonvulsant Screening Program (12).

13. Program, A. S. Ethosuximide Test 13 Results - Hippocampal Kindled Rats Anticonvulsant Screening Program (12).

14. Program, A. S. Ethosuximide Test 10 Results - Anticonvulsant Quantification (Rats I.P.) Anticonvulsant Screening Program (12).

22. Program, E. T. S. Gabapentin MES (no tox) Test 4 Results - Mice I.P. (506008). (2016).

23. Dalby, N. O. & Nielsen, E. B. Comparison of the preclinical anticonvulsant profiles of tiagabine, lamotrigine, gabapentin and vigabatrin. *Epilepsy Res* **28**, 63–72 (1997).

24. Program, A. S. Gabapentin Test 10 Results - Anticonvulsant Quantification (Rats I.P.) Anticonvulsant Screening Program (129018).

25. Program, E. T. S. Gabapentin (MES + TOX) Test 10 Results - Rats I.P. (506008).

31. Program, A. S. Gabapentin Test 26 Results - Corneal Kindled Mouse Anticonvulsant Screening Program (129018).

32. Program, E. T. S. Gabapentin Test 7 Results - 6Hz TOX, Mice.

33. Program, E. T. S. Gabapentin Test 25 Results - LTG-Resistant Amygdala Kindled Rat * * * * * *(506008).

43. Program, A. S. Carbamazepine Test 1 Results - Mice I.P. Identification Anticonvulsant Screening Program (4).

44. Program, A. S. Carbamazepine Test 16 Results - Anticonvulsant Quantification (Frings Mice) Anticonvulsant Screening Program (4).

45. Program, A. S. Carbamazepine Test 72 Results - Pilocarpine-induced Status (Interventional), Rats - Time 30 Min Anticonvulsant Screening Program (4).

46. Program, A. S. Carbamazepine Test 10 Results - Anticonvulsant Quantification (Rats I.P.) Anticonvulsant Screening Program (4).

62. Program, E. T. S. Lacosamide Test 4 Results - Mice I.P. (504009).

63. Program, E. T. S. Lacosamide Test 26 Results - Corneal Kindled Mouse (504009).

69. Program, A. S. Oxcarbazepine Test 1 Results - Mice I.P. Identification Anticonvulsant Screening Program (44001).

70. Program, A. S. Oxcarbazepine Test 4 Results - Mice I.P. Quantification Anticonvulsant Screening Program (44001).

71. Program, A. S. Oxcarbazepine Test 8 Results - Anticonvulsant Identification (Rats I.P.) Anticonvulsant Screening Program (339048).

72. Program, A. S. Oxcarbazepine Test 3 Results - Rat P.O. Quantification Anticonvulsant Screening Program (339048).

81. Program, A. S. Phenytoin Test 16 Results - Anticonvulsant Quantification (Frings Mice) Anticonvulsant Screening Program (1).

82. Program, A. S. Phenytoin Test 4 Results - Mice I.P. Quantification Anticonvulsant Screening Program (1).

83. Velíšek, L., Velíšková, J., Ptachewich, Y., Shinnar, S. & Moshé, S. L. Effects of MK‐801 and Phenytoin on Flurothyl‐Induced Seizures During Development. *Epilepsia* **36**, 179–185 (1995).

84. Program, E. T. S. Phenytoin Test 26 Results - Corneal Kindled Mouse (504005).

85. Rowley, N. M. & White, H. S. Comparative anticonvulsant efficacy in the corneal kindled mouse model of partial epilepsy: Correlation with other seizure and epilepsy models. *Epilepsy Res* **92**, 163–169 (2010).

86. Lotfy, D. M., Safar, M. M., Hassan, S. H. M. & Kenawy, S. A. Modulation of PTZ-induced convulsions in rats using topiramate alone or combined with low dose gamma irradiation: involving AKT/m-TOR pathway. *Toxicol. Mech. Methods* **32**, 18–26 (2022).

87. Pieróg, M. *et al.* Effects of new antiseizure drugs on seizure activity and anxiety-like behavior in adult zebrafish. *Toxicol. Appl. Pharmacol.* **427**, 115655 (2021).

88. Bialer, M., Twyman, R. E. & White, H. S. Correlation analysis between anticonvulsant ED50 values of antiepileptic drugs in mice and rats and their therapeutic doses and plasma levels. *Epilepsy Behav* **5**, 866–872 (2004).

89. Program, E. T. S. Topiramate Test 4 Results - Mice I.P. (503006).

90. Rigoulot, M., Boehrer, A. & Nehlig, A. Effects of Topiramate in Two Models of Genetically Determined Generalized Epilepsy, the GAERS and the Audiogenic Wistar AS. *Epilepsia* **44**, 14–19 (2003).

91. Wauquier, A. & Zhou, S. Topiramate: a potent anticonvulsant in the amygdala-kindled rat. *Epilepsy Res.* **24**, 73–77 (1996).

92. Program, A. S. Valproate Test 16 Results - Anticonvulsant Quantification (Frings Mice) Anticonvulsant Screening Program (8).

93. Program, A. S. Valproate Test 4 Results - Mice I.P. Quantification Anticonvulsant Screening Program (8).

94. Shenoy, A. K., Miyahara, J. T., Swinyard, E. A. & Kupferberg, H. J. Comparative Anticonvulsant Activity and Neurotoxicity of Clobazam, Diazepam, Phenobarbital, and Valproate in Mice and Rats. *Epilepsia* **23**, 399–408 (1982).

95. Program, A. S. Valproate Test 72 Results - Pilocarpine-induced Status, Rats - Time 30 Min Anticonvulsant Screening Program (8).

96. Program, A. S. Valproate Test 10 Results - Anticonvulsant Quantification (Rats I.P.) Anticonvulsant Screening Program (8).

97. Florek-Luszczki, M., Zagaja, M. & Luszczki, J. J. Influence of WIN 55,212-2 on the anticonvulsant and acute neurotoxic potential of clobazam and lacosamide in the maximal electroshock-induced seizure model and chimney test in mice. *Epilepsy Res.* **108**, 1728–1733 (2014).

98. Gatta, E. *et al.* Anticonvulsive Activity in Audiogenic DBA/2 Mice of 1,4-Benzodiazepines and 1,5-Benzodiazepines with Different Activities at Cerebellar Granule Cell GABAA Receptors. *J Mol Neurosci* **60**, 539–547 (2016).

99. Program, E. T. S. Clobazam Test 26 Results - Corneal Kindled Mouse (504003).

100. Program, E. T. S. Clobazam Test 4 Results - Mice I.P.

101. ICHIMARU, Y., GOMITA, Y. & MORIYAMA, M. EFFECTS OF CLOBAZAM ON AMYGDALOID AND HIPPOCAMPAL KINDLED SEIZURES IN RATS. *J. Pharmacobio-Dyn.* **10**, 189–194 (1987).

102. Program, E. T. S. Clobazam Test 10 Results - Rats I.P. (504003).

103. Tietz, E. I., Rosenberg, H. C. & Chiu, T. H. A comparison of the anticonvulsant effects of 1,4- and 1,5-benzodiazepines in the amygdala-kindled rat and their effects on motor function. *Epilepsy Res.* **3**, 31–40 (1989).

104. Program, A. S. Clonazepam Test 4 Results - Mice I.P. Quantification Anticonvulsant Screening Program (21).

105. Luszczki, J. J., Ratnaraj, N., Patsalos, P. N. & Czuczwar, S. J. Characterization of the Anticonvulsant, Behavioral and Pharmacokinetic Interaction Profiles of Stiripentol in Combination with Clonazepam, Ethosuximide, Phenobarbital, and Valproate Using Isobolographic Analysis. *Epilepsia* **47**, 1841–1854 (2006).

106. Program, A. S. Clonazepam Test 16 Results - Anticonvulsant Quantification (Frings Mice) Anticonvulsant Screening Program (21).

107. Program, A. S. Clonazepam Test 6 Results - BIC, PIC (Mice I.P.) Anticonvulsant Screening Program (21).

108. Program, A. S. Clonazepam Test 72 Results - Pilocarpine-induced Status, Rats - Time 30 Min Anticonvulsant Screening Program (21).

109. Program, A. S. Clonazepam Test 10 Results - Anticonvulsant Quantification (Rats I.P.) Anticonvulsant Screening Program (21).

110. Program, E. T. S. Clonazepam Test 26 Results - Corneal Kindled Mouse (504004).

111. Program, A. S. Clonazepam Test 13 Results - Hippocampal Kindled Rats Anticonvulsant Screening Program (21).

112. Program, A. S. Diazepam Test 4 Results - Mice I.P. Quantification Anticonvulsant Screening Program (7).

113. Program, A. S. Diazepam Test 71 Results - Pilocarpine-induced Status, Rats - Time 0 Min Anticonvulsant Screening Program (7).

114. Program, A. S. Diazepam Test 3 Results - Rat P.O. Quantification Anticonvulsant Screening Program (7).

115. Hanada, T., Ido, K. & Kosasa, T. Effect of perampanel, a novel AMPA antagonist, on benzodiazepine‐resistant status epilepticus in a lithium‐pilocarpine rat model. *Pharmacol. Res. Perspect.* **2**, e00063 (2014).

116. Hönack, D. & Löscher, W. Kindling Increases the Sensitivity of Rats to Adverse Effects of Certain Antiepileptic Drugs. *Epilepsia* **36**, 763–771 (1995).

117. Swinyard, E. A., Sofia, R. D. & Kupferberg, H. J. Comparative Anticonvulsant Activity and Neurotoxicity of Felbamate and Four Prototype Antiepileptic Drugs in Mice and Rats. *Epilepsia* **27**, 27–34 (1986).

118. Program, A. S. Felbamate Test 16 Results - Anticonvulsant Quantification (Frings Mice) Anticonvulsant Screening Program (3055).

119. Program, A. S. Felbamate Test 4 Results - Mice I.P. Quantification Anticonvulsant Screening Program (3055).

120. Sofia, R. D., Gordon, R., Gels, M. & Diamantis, W. Effects of felbamate and other anticonvulsant drugs in two models of status epilepticus in the rat. *Res. Commun. Chem. Pathol. Pharmacol.* **79**, 335–41 (1993).

121. Frey, H.-H. & Bartels, I. Felbamate and meprobamate: a comparison of their anticonvulsant properties. *Epilepsy Res.* **27**, 151–164 (1997).

122. Wlaź, P. & Löscher, W. Anticonvulsant Activity of Felbamate in Amygdala Kindling Model of Temporal Lobe Epilepsy in Rats. *Epilepsia* **38**, 1167–1172 (1997).

123. Program, A. S. Felbamate Test 10 Results - Anticonvulsant Quantification (Rats I.P.) Anticonvulsant Screening Program (3055).

129. Program, A. S. Phenobarbital Test 4 Results - Mice I.P. Quantification Anticonvulsant Screening Program (2).

130. Velíšek, L. *et al.* Age‐Dependent Effects of γ‐Aminobutyric Acid Agents on Flurothyl Seizures. *Epilepsia* **36**, 636–643 (1995).

131. Klitgaard, H., Matagne, A., Gobert, J. & Wülfert, E. Evidence for a unique profile of levetiracetam in rodent models of seizures and epilepsy. *Eur J Pharmacol* **353**, 191–206 (1998).

132. Program, A. S. Phenobarbital Test 72 Results - Pilocarpine-induced Status, Rats - Time 30 Min Anticonvulsant Screening Program (2).

133. Program, A. S. Phenobarbital Test 10 Results - Anticonvulsant Quantification (Rats I.P.) Anticonvulsant Screening Program (2).

134. Micheletti, G. *et al.* Antiepileptic drug evaluation in a new animal model: spontaneous petit mal epilepsy in the rat. *Arzneim.-Forsch.* **35**, 483–5 (1985).

135. Löscher, W. & Hönack, D. Comparison of the anticonvulsant efficacy of primidone and phenobarbital during chronic treatment of amygdala-kindled rats. *Eur. J. Pharmacol.* **162**, 309–322 (1989).

136. Sills, G. J., Butler, E., Thompson, G. G. & Brodie, M. J. Pharmacodynamic interaction studies with topiramate in the pentylenetetrazol and maximal electroshock seizure models. *Seizure* **13**, 287–295 (2004).

137. Program, A. S. Primidone Test 3 Results - Rat P.O. Quantification Anticonvulsant Screening Program (11).

146. Program, E. T. S. Tiagabine Test 7 Results - 6Hz, Mice (504007).

147. Program, E. T. S. Tiagabine Test 4 Results - Mice I.P. (504007).

148. Program, E. T. S. Tiagabine Test 26 Results - Corneal Kindled Mouse (504007).
