## Supplemental Figure 2 for "PAC – A novel translational concordance framework identifies preclinical seizure models with highest predictive validity for clinical focal onset seizures"

**Supplemental Figure 2. Data Sources for Preclinical ASM Response.** The preclinical efficacy of 32 FDA-approved ASMs currently available in the US was examined in a total of 23 seizure models across multiple species. Colors denote grading of *preclinical ASM response* based on reported TD_50_ and ED_50_ values for each model to calculate a protective index (PI), resulting in a weighted scale capturing relative preclinical anticonvulsant potential. Preclinical seizure models were grouped according to class/mechanism of action: calcium and sodium channel blockers, multimodal agents, GABAergic agents as well as agents with other mechanisms of action (including mTOR inhibitors, modulators of SV2A, selective serotonin reuptake inhibitors, and AMPA inhibitors). Individual data sources evaluated to assign depicted gradings are noted. m, mouse; r, rat; z, zebrafish; MES, maximal electroshock seizure, PTZ, pentylenetetrazole; preventative, treatment administered before pilocarpine-induced *status epilepticus*; interventional, treatment administered after pilocarpine-induced *status epilepticus*.

|  |  | **Generalized Seizure Models** | | | | | | | | | | | | **Focal Seizure Models** | | | | | | | | | | |
| --- | --- | --- | --- | --- | --- | --- | --- | --- | --- | --- | --- | --- | --- | --- | --- | --- | --- | --- | --- | --- | --- | --- | --- | --- |
|  |  | **MES** | | **scPTZ** | | | **Audiogenic** | **Bicuculline** | **Flurothyl** | **Picrotoxin** | ***Status* (Pilocarpine)** | | **GAERS or WAG/Rij** | **6-Hz (32 mA)** | | **6-Hz (44 mA)** | | **Systemic Kainate** | **Hippocampal Kainate** | **Amygdala Kainate** | **Corneal Kindling** | **Hippocampal Kindling** | **Amygdala Kindling** | **LTG-resistant** |
|  |  | **m** | **r** | **m** | **r** | **z** | **m** | **m** | **rodents** | **m** | **pre** | **int** | **r** | **m** | **r** | **m** | **r** | **r** | **m** | **m** | **m** | **r** | **r** | **r** |
| Ca_v_  channels | Ethosuximide | ^1^ | ^2^ | ^3^ | ^4^ | ^5^ | ^6,7^ | ^8^ | ^9^ | ^4^ | ^10^ |  | ^11,12^ | ^2^ | ^2^ | ^2^ | ^2^ | ^1^ |  |  |  | ^13,14^ | ^15^ | ^11^ |
|  | Methsuximide | ^16^ |  | ^16^ |  |  |  |  |  |  |  |  |  |  |  |  |  |  |  |  |  |  |  |  |
|  | Zonisamide | ^1^ | ^17^ | ^3^ | ^17^ | ^18^ |  | ^3^ | ^9^ | ^3^ |  |  | ^17,19^ | ^20^ |  |  |  | ^1^ |  |  |  |  | ^21^ |  |
|  | Gabapentin | ^22^ | ^2^ | ^23^ | ^24,25^ | ^26^ | ^23^ |  | ^2,27^ |  | ^11,28^ |  | ^29^ | ^2^ | ^2^ | ^2^ | ^2^ |  |  |  | ^25,30^ |  | ^23^ | ^25,31^ |
|  | Pregabalin | ^32^ | ^33^ | ^32^ | ^33^ | ^26^ | ^32^ |  |  |  | ^33,34^ | ^35^ | ^32^ | ^36,37^ |  |  |  |  | ^38^ |  |  | ^32^ | ^33^ |  |
| Na_v_  channels | Carbamazepine | ^39^ | ^39^ | ^3^ | ^40^ | ^26^ | ^41,42^ | ^39^ | ^9^ | ^39^ | ^10^ | ^43,44^ | ^29^ | ^2^ | ^2^ | ^2^ | ^2^ | ^11,45^ | ^38^ | ^46^ | ^47^ | ^39^ | ^48^ | ^11^ |
|  | Cenobamate | ^39^ | ^39^ | ^39^ | ^49^ |  | ^50^ | ^39^ |  | ^39^ | ^39^ |  | ^51^ | ^39^ |  | ^39^ |  |  |  |  |  | ^39^ |  |  |
|  | Eslicarbazepine Acetate | ^52^ | ^2^ |  |  |  |  |  |  |  |  |  |  | ^2^ | ^2^ | ^2^ | ^2^ |  |  |  | ^53^ |  |  | ^11^ |
|  | Fosphenytoin | ^54^ |  |  |  |  |  |  |  |  |  | ^55^ |  |  |  |  |  |  |  |  |  |  | ^56^ |  |
|  | Lacosamide | ^39^ | ^39^ | ^39^ | ^33^ | ^26^ | ^57^ | ^39^ |  | ^39^ |  | ^39,58^ | ^59^ | ^2^ | ^2^ | ^2^ | ^2^ |  |  |  | ^60,61^ | ^39^ | ^33^ | ^11^ |
|  | Lamotrigine | ^62^ | ^39^ | ^39^ | ^63^ | ^18^ | ^23^ | ^39^ | ^2,64^ | ^39^ |  |  | ^65^ | ^2^ | ^2^ | ^2^ | ^2^ | ^66^ | ^38^ |  | ^47^ | ^39^ | ^11,48^ | ^11^ |
|  | Oxcarbazepine | ^67,68^ | ^69,70^ | ^71^ |  | ^5^ | ^72,73^ |  |  |  |  |  | ^74^ | ^73,75^ |  |  |  | ^69,70,76^ |  |  |  |  | ^77^ |  |
|  | Phenytoin | ^39^ | ^2^ | ^3^ | ^4^ | ^78^ | ^79,80^ | ^39^ | ^81^ | ^39^ | ^10^ | ^55^ | ^12^ | ^2^ | ^2^ | ^2^ | ^2^ | ^1^ |  | ^46^ | ^80,82^ | ^39^ | ^48^ | ^11^ |
|  | Rufinamide | ^4^ | ^2^ | ^4^ | ^4^ |  |  | ^4^ |  | ^4^ |  |  |  | ^2^ | ^2^ | ^2^ | ^2^ |  |  |  |  |  |  | ^11^ |
| Multimodal | Topiramate | ^39^ | ^2^ | ^83^ | ^11,84^ | ^85^ | ^86,87^ | ^39^ |  | ^39^ |  | ^39^ | ^39,88^ | ^2^ | ^2^ | ^2^ | ^2^ | ^66^ |  |  | ^30^ | ^39^ | ^11,89^ | ^11^ |
|  | Valproate | ^8^ | ^2^ | ^8^ | ^8^ | ^5^ | ^90,91^ | ^92^ | ^2,64^ | ^39^ | ^10^ | ^93,94^ | ^39,88^ | ^2^ | ^2^ | ^2^ | ^2^ | ^1^ | ^38^ | ^46^ | ^47^ | ^39^ | ^33^ | ^11^ |
| GABA | Clobazam | ^95^ | ^92^ | ^92^ | ^92^ |  | ^95,96^ | ^92^ |  | ^92^ |  |  |  | ^2^ | ^2^ | ^2^ | ^2^ |  |  |  | ^97,98^ | ^99,100^ | ^101^ | ^11^ |
|  | Clonazepam | ^102^ | ^2^ | ^103^ | ^40^ | ^78^ | ^102,104^ | ^102,105^ | ^9^ | ^102,105^ | ^10,40^ | ^106,107^ |  | ^2^ | ^2^ | ^2^ | ^2^ | ^45^ |  |  | ^102,108^ | ^107,109^ | ^101^ | ^11^ |
|  | Diazepam | ^92^ | ^92^ | ^3^ | ^92^ | ^26^ | ^96,110^ | ^92^ | ^9^ | ^92^ | ^111,112^ | ^113^ | ^12,92^ | ^47^ |  | ^47^ |  | ^1^ | ^38^ | ^46^ | ^47^ | ^99,114^ | ^101^ |  |
|  | Felbamate | ^39^ | ^39^ | ^115^ | ^115^ | ^85^ | ^116,117^ | ^39^ |  | ^39^ | ^39,118^ |  | ^39,119^ | ^39^ |  | ^39^ |  |  |  |  |  | ^39^ | ^120,121^ |  |
|  | Ganaxolone | ^8^ | ^8^ | ^8^ | ^8^ |  |  | ^8^ | ^8,122^ |  |  | ^123^ |  | ^8,124^ |  |  |  |  |  |  |  |  |  |  |
|  | Phenobarbital | ^92^ | ^2^ | ^92^ | ^92^ | ^125^ | ^126,127^ | ^92^ | ^128^ | ^92^ | ^129^ | ^130,131^ | ^132^ | ^2,39,47^ | ^2^ | ^2^ | ^2^ | ^11,45^ | ^38^ | ^46^ | ^47^ | ^39^ | ^11,133^ | ^11^ |
|  | Primidone | ^40,134^ | ^135^ | ^136^ | ^40^ |  | ^40,127^ | ^136^ |  |  |  |  |  |  |  |  |  |  |  |  |  |  | ^133^ |  |
|  | Stiripentol | ^137,138^ |  | ^103^ | ^139,140^ |  |  |  |  |  | ^140^ |  | ^139,141^ |  |  |  |  |  |  |  |  |  |  |  |
| Other/ Unknown | Tiagabine | ^142^ | ^2^ | ^23^ | ^143^ | ^5^ | ^23^ |  |  |  |  |  | ^29^ | ^2^ | ^2^ | ^2^ | ^2^ |  | ^38,144^ |  | ^145,146^ | ^11,83^ | ^23^ | ^11^ |
|  | Vigabatrin | ^142^ |  | ^23,147^ | ^143^ |  | ^23^ |  | ^128^ | ^23,148^ | ^23,28^ |  | ^149^ | ^150^ |  |  |  | ^23,151^ | ^38^ |  | ^83^ |  | ^23^ |  |
|  | Brivaracetam | ^152^ |  | ^153^ |  |  | ^153^ | ^153^ |  |  | ^154^ |  | ^155^ | ^153^ |  |  |  |  | ^156^ |  | ^155^ | ^155^ | ^155^ |  |
|  | Levetiracetam | ^152^ | ^2^ | ^3^ | ^33^ | ^18^ | ^153^ | ^153^ |  | ^3^ | ^129^ | ^157^ | ^155^ | ^47^ | ^2^ | ^47^ | ^2^ | ^1^ | ^38^ |  | ^47^ | ^155^ | ^155^ | ^11^ |
|  | Everolimus |  |  |  |  |  |  |  |  |  |  |  |  |  |  |  |  |  | ^156^ |  |  |  |  |  |
|  | Fenfluramine | ^158^ | ^159^ | ^160^ | ^160^ |  | ^159,161^ |  |  |  |  |  |  | ^160^ |  | ^159^ |  |  |  |  |  |  |  |  |
|  | Perampanel | ^162^ |  | ^162^ | ^163,164^ | ^18^ | ^164^ |  |  |  |  | ^113^ | ^165^ | ^164^ |  | ^164^ |  |  | ^156,164^ |  | ^30,164^ |  | ^21^ |  |
|  | Cannabidiol | ^166^ | ^167^ | ^167^ | ^168^ | ^169^ | ^167,170^ | ^167,171^ |  | ^167,171^ | ^172^ |  | ^173^ | ^172^ |  | ^167^ |  | ^172,174^ |  |  | ^167^ |  | ^172,175^ | ^167^ |

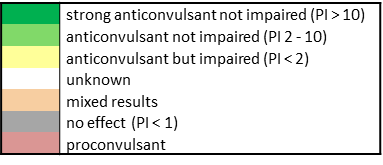

6. Ethosuximide (ETSP #12) - Frings Mice (Test 16), National Institute of Neurological Disorders and Stroke Epilepsy Therapy Screening Program, PANAChE Database.

7. Ethosuximide (ETSP #12) - MES, scPTZ, Mice I.P. (Test 4), National Institute of Neurological Disorders and Stroke Epilepsy Therapy Screening Program, PANAChE Database.

13. Ethosuximide (ETSP #12) - Hippocampal Kindled Rats (Test 13), National Institute of Neurological Disorders and Stroke Epilepsy Therapy Screening Program, PANAChE Database.

14. Ethosuximide (ETSP #12) - MES, scPTZ, Rats I.P. (Test 10), National Institute of Neurological Disorders and Stroke Epilepsy Therapy Screening Program, PANAChE Database.

22. Gabapentin (ETSP #506008) - MES, scPTZ, Mice I.P. (Test 4), National Institute of Neurological Disorders and Stroke Epilepsy Therapy Screening Program, PANAChE Database

23. Dalby, N. O. & Nielsen, E. B. Comparison of the preclinical anticonvulsant profiles of tiagabine, lamotrigine, gabapentin and vigabatrin. *Epilepsy Res* **28**, 63–72 (1997).

24. Gabapentin (ETSP #129018) - MES, scPTZ, Rats I.P. (Test 10), National Institute of Neurological Disorders and Stroke Epilepsy Therapy Screening Program, PANAChE Database.

25. Gabapentin (ETSP #506008) - MES, scPTZ, Rats I.P. (Test 10), National Institute of Neurological Disorders and Stroke Epilepsy Therapy Screening Program, PANAChE Database.

31. Gabapentin (ETSP #506008) - LTG-Resistant Amygdala Kindled Rat (Test 25), National Institute of Neurological Disorders and Stroke Epilepsy Therapy Screening Program, PANAChE Database.

41. Carbamazepine (ETSP #4) - MES, scPTZ, TOX, Mice I.P. (Test 1), National Institute of Neurological Disorders and Stroke Epilepsy Therapy Screening Program, PANAChE Database.

42. Carbamazepine (ETSP #4) - Frings Mice (Test 16), National Institute of Neurological Disorders and Stroke Epilepsy Therapy Screening Program, PANAChE Database.

43. Carbamazepine (ETSP #4) - Pilocarpine-induced Status Epilepticus- Acute Intervention, Rats (Test 72), National Institute of Neurological Disorders and Stroke Epilepsy Therapy Screening Program, PANAChE Database.

44. Carbamazepine (ETSP #4) - MES, scPTZ, Rats I.P. (Test 10), National Institute of Neurological Disorders and Stroke Epilepsy Therapy Screening Program, PANAChE Database.

60. Lacosamide (ETSP #504009) - MES, scPTZ, Mice I.P. (Test 4), National Institute of Neurological Disorders and Stroke Epilepsy Therapy Screening Program, PANAChE Database.

61. Lacosamide (ETSP #504009) - Corneal Kindled Mouse (Test 26), National Institute of Neurological Disorders and Stroke Epilepsy Therapy Screening Program, PANAChE Database.

67. Oxcarbazepine (ETSP #44001) - MES, scPTZ, TOX, Mice I.P. (Test 1), National Institute of Neurological Disorders and Stroke Epilepsy Therapy Screening Program, PANAChE Database.

68. Oxcarbazepine (ETSP #44001) - MES, scPTZ, TOX, Mice I.P. (Test 4), National Institute of Neurological Disorders and Stroke Epilepsy Therapy Screening Program, PANAChE Database.

69. Oxcarbazepine (ETSP #339048) - MES, scPTZ, Rats I.P. (Test 8), National Institute of Neurological Disorders and Stroke Epilepsy Therapy Screening Program, PANAChE Database.

70. Oxcarbazepine (ETSP #339048) - MES, scPTZ, Rats P.O. (Test 3), National Institute of Neurological Disorders and Stroke Epilepsy Therapy Screening Program, PANAChE Database.

79. Phenytoin (ETSP #1) - Anticonvulsant Quantification, Frings Mice, Mice I.P. (Test 16), National Institute of Neurological Disorders and Stroke Epilepsy Therapy Screening Program, PANAChE Database.

80. Phenytoin (ETSP #1) - MES, scPTZ, Mice I.P. (Test 4), National Institute of Neurological Disorders and Stroke Epilepsy Therapy Screening Program, PANAChE Database.

81. Velíšek, L., Velíšková, J., Ptachewich, Y., Shinnar, S. & Moshé, S. L. Effects of MK‐801 and Phenytoin on Flurothyl‐Induced Seizures During Development. *Epilepsia* **36**, 179–185 (1995).

82. Program, E. T. S. Phenytoin Test 26 Results - Corneal Kindled Mouse (504005).

83. Rowley, N. M. & White, H. S. Comparative anticonvulsant efficacy in the corneal kindled mouse model of partial epilepsy: Correlation with other seizure and epilepsy models. *Epilepsy Res* **92**, 163–169 (2010).

84. Lotfy, D. M., Safar, M. M., Hassan, S. H. M. & Kenawy, S. A. Modulation of PTZ-induced convulsions in rats using topiramate alone or combined with low dose gamma irradiation: involving AKT/m-TOR pathway. *Toxicol. Mech. Methods* **32**, 18–26 (2022).

85. Pieróg, M. *et al.* Effects of new antiseizure drugs on seizure activity and anxiety-like behavior in adult zebrafish. *Toxicol. Appl. Pharmacol.* **427**, 115655 (2021).

86. Bialer, M., Twyman, R. E. & White, H. S. Correlation analysis between anticonvulsant ED50 values of antiepileptic drugs in mice and rats and their therapeutic doses and plasma levels. *Epilepsy Behav* **5**, 866–872 (2004).

87. Topiramate (ETSP #503006) - MES, scPTZ, Mice I.P. (Test 4), National Institute of Neurological Disorders and Stroke Epilepsy Therapy Screening Program, PANAChE Database.

88. Rigoulot, M., Boehrer, A. & Nehlig, A. Effects of Topiramate in Two Models of Genetically Determined Generalized Epilepsy, the GAERS and the Audiogenic Wistar AS. *Epilepsia* **44**, 14–19 (2003).

89. Wauquier, A. & Zhou, S. Topiramate: a potent anticonvulsant in the amygdala-kindled rat. *Epilepsy Res.* **24**, 73–77 (1996).

90. Valproate (ETSP #8) - Frings Mice (Test 16), National Institute of Neurological Disorders and Stroke Epilepsy Therapy Screening Program, PANAChE Database.

91. Valproate (ETSP #8) - MES, scPTZ, Mice I.P. (Test 4), National Institute of Neurological Disorders and Stroke Epilepsy Therapy Screening Program, PANAChE Database.

92. Shenoy, A. K., Miyahara, J. T., Swinyard, E. A. & Kupferberg, H. J. Comparative Anticonvulsant Activity and Neurotoxicity of Clobazam, Diazepam, Phenobarbital, and Valproate in Mice and Rats. *Epilepsia* **23**, 399–408 (1982).

93. Valproate (ETSP #8) - Pilocarpine-induced Status Epilepticus- Acute Intervention, Rats (Test 72), National Institute of Neurological Disorders and Stroke Epilepsy Therapy Screening Program, PANAChE Database.

94. Valproate (ETSP #8) - MES, scPTZ, Rats I.P. (Test 10), National Institute of Neurological Disorders and Stroke Epilepsy Therapy Screening Program, PANAChE Database.

95. Florek-Luszczki, M., Zagaja, M. & Luszczki, J. J. Influence of WIN 55,212-2 on the anticonvulsant and acute neurotoxic potential of clobazam and lacosamide in the maximal electroshock-induced seizure model and chimney test in mice. *Epilepsy Res.* **108**, 1728–1733 (2014).

96. Gatta, E. *et al.* Anticonvulsive Activity in Audiogenic DBA/2 Mice of 1,4-Benzodiazepines and 1,5-Benzodiazepines with Different Activities at Cerebellar Granule Cell GABAA Receptors. *J Mol Neurosci* **60**, 539–547 (2016).

97. Clobazam (ETSP #504003) - Corneal Kindled Mouse (Test 26), National Institute of Neurological Disorders and Stroke Epilepsy Therapy Screening Program, PANAChE Database.

98. Clobazam (ETSP #504003) - MES, scPTZ, Mice I.P. (Test 4), National Institute of Neurological Disorders and Stroke Epilepsy Therapy Screening Program, PANAChE Database.

99. ICHIMARU, Y., GOMITA, Y. & MORIYAMA, M. EFFECTS OF CLOBAZAM ON AMYGDALOID AND HIPPOCAMPAL KINDLED SEIZURES IN RATS. *J. Pharmacobio-Dyn.* **10**, 189–194 (1987).

100. Clobazam (ETSP #504003) - MES, scPTZ, Rats I.P. (Test 10), National Institute of Neurological Disorders and Stroke Epilepsy Therapy Screening Program, PANAChE Database.

101. Tietz, E. I., Rosenberg, H. C. & Chiu, T. H. A comparison of the anticonvulsant effects of 1,4- and 1,5-benzodiazepines in the amygdala-kindled rat and their effects on motor function. *Epilepsy Res.* **3**, 31–40 (1989).

102. Clonazepam (ETSP #21) - MES, scPTZ, Mice I.P. (Test 4), National Institute of Neurological Disorders and Stroke Epilepsy Therapy Screening Program, PANAChE Database.

103. Luszczki, J. J., Ratnaraj, N., Patsalos, P. N. & Czuczwar, S. J. Characterization of the Anticonvulsant, Behavioral and Pharmacokinetic Interaction Profiles of Stiripentol in Combination with Clonazepam, Ethosuximide, Phenobarbital, and Valproate Using Isobolographic Analysis. *Epilepsia* **47**, 1841–1854 (2006).

104. Clonazepam (ETSP #21) - Frings Mice (Test 16), National Institute of Neurological Disorders and Stroke Epilepsy Therapy Screening Program, PANAChE Database.

105. Clonazepam (ETSP #21) - BIC, PIC, Mice I.P. (Test 6), National Institute of Neurological Disorders and Stroke Epilepsy Therapy Screening Program, PANAChE Database.

106. Clonazepam (ETSP #21) - Pilocarpine-induced Status Epilepticus- Acute Intervention, Rats (Test 72), National Institute of Neurological Disorders and Stroke Epilepsy Therapy Screening Program, PANAChE Database.

107. Clonazepam (ETSP #21) - MES, scPTZ, Rats I.P. (Test 10), National Institute of Neurological Disorders and Stroke Epilepsy Therapy Screening Program, PANAChE Database.

108. Clonazepam (ETSP #504004) - Corneal Kindled Mouse (Test 26), National Institute of Neurological Disorders and Stroke Epilepsy Therapy Screening Program, PANAChE Database.

109. Clonazepam (ETSP #21) - Hippocampal Kindled Rats (Test 13), National Institute of Neurological Disorders and Stroke Epilepsy Therapy Screening Program, PANAChE Database.

110. Diazepam (ETSP #7) - MES, scPTZ, Mice I.P. (Test 4), National Institute of Neurological Disorders and Stroke Epilepsy Therapy Screening Program, PANAChE Database.

111. Diazepam (ETSP #7) - Pilocarpine-induced Status Epilepticus, Rats (Test 71), National Institute of Neurological Disorders and Stroke Epilepsy Therapy Screening Program, PANAChE Database.

112. Diazepam (ETSP #7) - MES, scPTZ, Rats P.O. (Test 3), National Institute of Neurological Disorders and Stroke Epilepsy Therapy Screening Program, PANAChE Database.

113. Hanada, T., Ido, K. & Kosasa, T. Effect of perampanel, a novel AMPA antagonist, on benzodiazepine‐resistant status epilepticus in a lithium‐pilocarpine rat model. *Pharmacol. Res. Perspect.* **2**, e00063 (2014).

114. Hönack, D. & Löscher, W. Kindling Increases the Sensitivity of Rats to Adverse Effects of Certain Antiepileptic Drugs. *Epilepsia* **36**, 763–771 (1995).

115. Swinyard, E. A., Sofia, R. D. & Kupferberg, H. J. Comparative Anticonvulsant Activity and Neurotoxicity of Felbamate and Four Prototype Antiepileptic Drugs in Mice and Rats. *Epilepsia* **27**, 27–34 (1986).

116. Felbamate (ETSP #3055) - Frings Mice (Test 16), National Institute of Neurological Disorders and Stroke Epilepsy Therapy Screening Program, PANAChE Database.

117. Felbamate (ETSP #3055) - MES, scPTZ, Mice I.P. (Test 4), National Institute of Neurological Disorders and Stroke Epilepsy Therapy Screening Program, PANAChE Database.

118. Sofia, R. D., Gordon, R., Gels, M. & Diamantis, W. Effects of felbamate and other anticonvulsant drugs in two models of status epilepticus in the rat. *Res. Commun. Chem. Pathol. Pharmacol.* **79**, 335–41 (1993).

119. Frey, H.-H. & Bartels, I. Felbamate and meprobamate: a comparison of their anticonvulsant properties. *Epilepsy Res.* **27**, 151–164 (1997).

120. Felbamate (ETSP #3055) - MES, scPTZ, Rats I.P. (Test 10), National Institute of Neurological Disorders and Stroke Epilepsy Therapy Screening Program, PANAChE Database.

126. Phenobarbital (ETSP #2) - MES, scPTZ, Mice I.P. (Test 4), National Institute of Neurological Disorders and Stroke Epilepsy Therapy Screening Program, PANAChE Database.

127. COLLINS, A. J. & HORLINGTON, M. A sequential screening test based on the running component of audiogenic seizures in mice, including reference compound PD50 values. *Br. J. Pharmacol.* **37**, 140–150 (1969).

128. Velíšek, L. *et al.* Age‐Dependent Effects of γ‐Aminobutyric Acid Agents on Flurothyl Seizures. *Epilepsia* **36**, 636–643 (1995).

129. Klitgaard, H., Matagne, A., Gobert, J. & Wülfert, E. Evidence for a unique profile of levetiracetam in rodent models of seizures and epilepsy. *Eur J Pharmacol* **353**, 191–206 (1998).

130. Phenobarbital (ETSP #2) - Pilocarpine-induced Status Epilepticus- Acute Intervention, Rats (Test 72), National Institute of Neurological Disorders and Stroke Epilepsy Therapy Screening Program, PANAChE Database.

131. Phenobarbital (ETSP #2) - MES, scPTZ, Rats I.P. (Test 10), National Institute of Neurological Disorders and Stroke Epilepsy Therapy Screening Program, PANAChE Database.

132. Micheletti, G. *et al.* Antiepileptic drug evaluation in a new animal model: spontaneous petit mal epilepsy in the rat. *Arzneim.-Forsch.* **35**, 483–5 (1985).

133. Löscher, W. & Hönack, D. Comparison of the anticonvulsant efficacy of primidone and phenobarbital during chronic treatment of amygdala-kindled rats. *Eur. J. Pharmacol.* **162**, 309–322 (1989).

134. Sills, G. J., Butler, E., Thompson, G. G. & Brodie, M. J. Pharmacodynamic interaction studies with topiramate in the pentylenetetrazol and maximal electroshock seizure models. *Seizure* **13**, 287–295 (2004).

135. Primidone (ETSP #11) -MES, scPTZ, Rats P.O. (Test 3), National Institute of Neurological Disorders and Stroke Epilepsy Therapy Screening Program, PANAChE Database.

144. Tiagabine (ETSP #504007) - 6Hz, Mice (Test 7), National Institute of Neurological Disorders and Stroke Epilepsy Therapy Screening Program, PANAChE Database.

145. Tiagabine (ETSP #504007) - MES, scPTZ, Mice I.P. (Test 4), National Institute of Neurological Disorders and Stroke Epilepsy Therapy Screening Program, PANAChE Database.

146. Tiagabine (ETSP #504007) - Corneal Kindled Mouse (Test 26), National Institute of Neurological Disorders and Stroke Epilepsy Therapy Screening Program, PANAChE Database.
