## Supplemental Figure 3 for "PAC – A novel translational concordance framework identifies preclinical seizure models with highest predictive validity for clinical focal onset seizures"

**Supplemental Figure 3. Data Sources for Clinical ASM Response.** Clinical efficacy of the 32 FDA-approved ASMs was evaluated based on established reports of perceived efficacy and use. Colors denote grading of clinical ASM response based on prescribing patterns for FOS, resulting in a weighted scale capturing relative clinical anticonvulsant potential. Individual data sources evaluated to assign depicted gradings are noted.

|  |  | **FOS** |
| --- | --- | --- |
| Ca_v_ channels | Ethosuximide | ^1,2^ |
|  | Methsuximide | ^3^ |
|  | Zonisamide | ^2^ |
|  | Gabapentin | ^1^ |
|  | Pregabalin | ^1^ |
| Na_v_ channels | Carbamazepine | ^1,2,4^ |
|  | Cenobamate | ^5^ |
|  | Eslicarbazepine Acetate | ^1^ |
|  | Fosphenytoin | ^6^ |
|  | Lacosamide | ^1,2,4^ |
|  | Lamotrigine | ^1,2,4^ |
|  | Oxcarbazepine | ^1,2,4^ |
|  | Phenytoin | ^2^ |
|  | Rufinamide | ^1^ |
| Multimodal | Topiramate | ^2^ |
|  | Valproate | ^1,2,4^ |
| GABA | Clobazam | ^1^ |
|  | Clonazepam | ^1,4^ |
|  | Diazepam | ^7^ |
|  | Felbamate | ^1,8^ |
|  | Ganaxolone | ^9^ |
|  | Phenobarbital | ^2^ |
|  | Primidone | ^1^ |
|  | Stiripentol | ^10^ |
|  | Tiagabine | ^1^ |
|  | Vigabatrin | ^1^ |
| Other/Unknown | Brivaracetam | ^1^ |
|  | Levetiracetam | ^1,4^ |
|  | Everolimus |  |
|  | Fenfluramine |  |
|  | Perampanel | ^11^ |
|  | Cannabidiol |  |

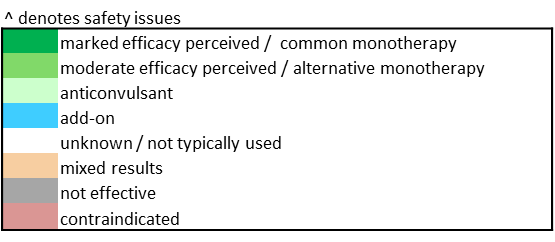
